# Single-kernel near-infrared spectroscopy enables haploid kernel sorting in field and sweet corn using high-oil haploid inducers across diverse donor-inducer combinations

**DOI:** 10.64898/2026.07.23.740370

**Authors:** Suman Sharma, Jeffery L. Gustin, Ursula K. Frei, A. Mark Settles, Thomas Lübberstedt, Márcio F. R. Resende, Jenna Hershberger

**Affiliations:** Plant and Environmental Sciences Department, Pee Dee Research and Education Center, Clemson University, Florence, SC, USA; Maize Genetics Cooperation Stock Center, USDA-ARS, Urbana, IL, USA; Horticultural Sciences Department, University of Florida, Gainesville, FL, USA; Department of Agronomy, Iowa State University, Ames, IA, USA; Space Biosciences Division, NASA Ames Research Center, Moffett Field, Mountain View, CA, 94035, USA

**Keywords:** Doubled haploid, corn, NIR spectroscopy, kernel oil content, support vector machine

## Abstract

**Key message:** A single-kernel near-infrared reflectance spectroscopy-based sorter can effectively identify haploid kernels for doubled haploid production in field and sweet corn backgrounds.

Doubled haploid (DH) technology significantly shortens the breeding cycle for developing homozygous inbred lines in maize (*Zea mays*). Manual sorting of haploids from a larger bulk of hybrid kernels in an induction cross is a major bottleneck in DH development. Automated systems based on near-infrared (NIR) reflectance spectroscopy can be valuable tools for rapid haploid sorting, provided that sorting accuracy is sufficient for incorporation into the DH process. In this study, we evaluated the accuracy of a custom-built single-kernel NIR (skNIR) sorter for classifying haploid kernels from 12 high-oil haploid induction populations generated from two sweet corn and two field corn donors and four high-oil haploid inducers (HOHIs). We evaluated several general classification models that can be applied without population-specific recalibration or prior genotyping, including models that classified haploids based solely on predicted oil content, as well as multivariate methods that used all wavelengths of the NIR spectra. The highest classification accuracy was obtained using a general multivariate support vector machine (SVM) model. When combined with the two best-performing HOHIs, the general SVM model accurately sorted induction populations from two of the three donor backgrounds crossed with these inducers. Two oil-based methods showed less accurate classification than the multivariate SVM model, due to overlapping oil content distributions across the two kernel classes. Overall, this study demonstrates effective skNIR-based sorting of haploid kernels from diverse induction populations using a single general model. The practical deployment of this instrument in maize breeding programs is discussed.

## 1. Introduction

The use of doubled haploid (DH) technology has expedited maize (*Zea mays*) breeding efforts through the efficient production of homozygous inbred lines (Geiger and Gordillo 2009; reviewed by Liu et al. 2016). DHs enable the development of new inbred lines in just two generations compared to traditional breeding methods that require six or more generations of self-pollination. This approach enables both field corn and sweet corn breeders to rapidly assess the hybrid performance of many inbred lines, significantly reducing the time required for variety release (Hallauer et al. 2010). DH technology also facilitates faster genetic improvement by shortening the breeding cycles required for the introgression of desirable alleles into elite maize germplasm (Lübberstedt and Frei 2012).

DHs in maize are produced through a four-step process: (1) haploid induction, (2) haploid identification, (3) genome doubling of confirmed haploids, and (4) self-pollination of DHs to produce fully homozygous lines (reviewed by Chaikam et al. 2019; reviewed by Dermail et al. 2024b). Haploids are induced *in vivo* by crossing a male haploid-inducer line with a female donor line. The frequency of haploid induction in maize is typically low, below 0.1% (Chase 1949). However, the discovery of Stock 6 (Coe 1959) with increased haploid induction frequency and subsequent improvement through breeding has resulted in modern maternal haploid inducer lines with induction frequencies up to 15% (Coe 1959; Cai et al. 2007; reviewed by Liu et al. 2016; reviewed by Trentin et al. 2020). Despite the increased haploid induction rate (HIR), haploids still represent only a small proportion of the total kernels in an induction cross, making haploid sorting a necessary but time-consuming step in the *in vivo* haploid induction process.

Identification of haploid kernels within an induction cross population is commonly performed visually using a kernel color marker (reviewed by Maqbool et al. 2020). This technique relies on a dominant color marker allele inherited from the male parent (paternal genome) (Nanda and Chase 1966; Geiger and Gordillo 2009; reviewed by Liu et al. 2016). The *R1-navajo* (*R1-nj*) allele of the *R1* gene induces purple coloration via anthocyanin accumulation in both the embryo and the aleurone layer of the endosperm and is often used for haploid identification. In a cross between a colorless kernel female donor and a male inducer carrying the *R1-nj* allele, haploids can be visually distinguished from diploid hybrid kernels by the absence of anthocyanin accumulation in the haploid embryo. While effective, this classical method is laborious and time-consuming and can be limited by suppression of kernel pigmentation in some genetic backgrounds (Chaikam and Prasanna 2012). Kernel color suppression poses a particular problem in sweet corn, where anthocyanin accumulation in both kernels and plants is inhibited to meet consumer preferences, limiting accurate haploid identification (Tracy 2001; Chaikam et al. 2015). A red root marker, governed by the dominant *purple plant 1* allele of the *Pl1* gene (Rotarenco et al. 2010), serves as an alternative color marker for haploid identification at the seedling stage. These visual markers can be subjective and require trained personnel to manually sort kernels or seedlings, which is a bottleneck in large-scale DH production.

Automated sorting systems are a potentially efficient alternative to manual haploid kernel sorting (reviewed by Dermail et al. 2024b). Image-based systems that rely on the *R1-nj* marker have been shown to sort haploid kernels with a reported accuracy of over 90% (Song et al. 2018). As with manual sorting, the effectiveness of these systems can be limited by the inconsistent penetrance of pigmentation markers across diverse genetic backgrounds, resulting in misclassification of true haploids as hybrids or requiring custom recalibration of prediction algorithms for some induction crosses. Chemical differences between kernel classes offer an alternative basis for automated classification. Kernel oil content is among the most reliable chemical markers for distinguishing between haploid and hybrid kernels (Rotarenco et al. 2007; Melchinger et al. 2013; Gustin et al. 2020). Kernel oil predominantly accumulates in the embryo and scutellum, with very little present in the starchy endosperm. In haploid kernels, oil content reflects only the contribution of the female parent, while hybrid kernels, which receive a male genetic contribution, tend to have higher oil content due to the xenia effect (Chen and Song 2003).

To maximize the difference in oil content between haploid and hybrid kernels, haploid inducer lines with a high kernel oil content have been developed (Melchinger et al. 2013, 2014; Dong et al. 2014; Liu et al. 2022). These high-oil haploid inducers (HOHIs) significantly increase the oil content of hybrid kernels, widening the relative difference in oil content between haploids and hybrids (Melchinger et al. 2013, 2014; Dong et al. 2014). Pairing HOHIs with nuclear magnetic resonance (NMR) spectroscopy enables automated haploid sorting across diverse induction populations based on relative oil content differences between kernel classes (Liu et al. 2012; Wang et al. 2016; Melchinger et al. 2017; Qu et al. 2021). The accuracy of NMR-based systems can exceed 95%, but this method is relatively slow due to the time required to weigh and measure each kernel in the NMR magnetic detector (Wang et al. 2016).

Near-infrared (NIR) spectroscopic techniques offer a fast and cost-effective solution for haploid kernel separation (reviewed by Rathna Priya and Manickavasagan 2021; reviewed by Dermail et al. 2024b). The single-kernel NIR (skNIR) system developed by Armstrong (2006) captures spectral data in a fraction of a second per kernel. Multiple kernel traits can be simultaneously predicted from the spectra, such as relative oil, protein, and starch content, in addition to kernel weight, volume, and density (Armstrong 2006; Armstrong and Tallada 2012; Spielbauer et al. 2009; Gustin et al. 2013). Gustin et al. (2020) demonstrated that the skNIR system could enrich the haploid pool from 12% haploids in the starting induction cross to above 50% haploids across multiple induction crosses using a conventional (non-high-oil) inducer. The most discriminating NIR-determined trait was relative oil content, with an average difference of 0.88% between haploid and hybrid kernels. Importantly, induction populations with an oil difference greater than 1.7% had high sorting accuracy. Haploid classification accuracies improved further when models used full skNIR spectra rather than skNIR-predicted oil content alone, indicating that skNIR classification models can extract more information than just the oil signal to separate haploid and hybrid kernels.

Here, we evaluated the sorting accuracy of a custom-built skNIR sorter for sorting haploid kernels by employing HOHIs for developing induction populations and further extended the approach to sweet corn. In addition to haploid and hybrid kernels, defective kernels from the induction populations were also included in the analysis to assess the sorting accuracy of the three naturally occurring kernel classes (Zhao et al. 2013; Kelliher et al. 2017; Liu et al. 2018; Qu et al. 2020). The current study includes 12 induction cross populations generated using field and sweet corn donors and HOHIs with the following objectives: (1) compare classification accuracies between visual sorting based on *R1-nj* marker, oil-based methods, and multivariate spectral methods for effective and rapid sorting of haploid kernels; and (2) investigate the influence of parental genetic backgrounds on the relative difference in kernel oil content between haploid and hybrid kernels generated from a high-oil inducer. Our goal is to identify haploid sorting approaches that are operationally effective, minimize classification error rates, and are broadly applicable across diverse backgrounds encountered in DH production.

## 2. Materials and Methods

### 2.1. Kernel samples

Kernels from 12 induction crosses were used in this study (**Table 1**). Four commercial hybrids were used as female parents (donors). Viking 51 and Viking 60 are field corn hybrids obtained from Albert Lea Seeds (Albert Lea, MN) that have strong and consistent expression of anthocyanin kernel color and are used by the Iowa State University Doubled Haploid Facility as testers for evaluating new haploid inducers. CCO-1 and CCO-2 are sweet corn hybrids donated by Crookham Company (Caldwell, ID) that have moderately suppressed anthocyanin accumulation in the kernel and represent a challenging genetic background for visual haploid identification. Four HOHIs were used as male parents (IHO-1 through IHO-4). The HOHIs were developed by crossing IHO, a high oil inbred line (∼17% oil content) developed from the Illinois long-term selection experiment (Moose et al. 2004), with BHI305, a high-quality haploid inducer used by the Iowa State University Doubled Haploid Facility (average HIR of 10-12%; Trentin et al. 2020). Marker-assisted backcrossing was used to select for the haploid induction QTL *qhir1* and *qhir8* (Prigge et al. 2012), and skNIR was used to select high-oil progeny ears to generate four HOHIs. The HOHIs contained the *R1-nj* and *Pl1* dominant anthocyanin markers, had HIRs ranging from 3% to 13%, and oil content between 9.6% and 11.4%. The induction crosses were made in 2021 at the Iowa State University Agricultural Engineering and Agronomy Research Farm in Boone, IA using controlled pollinations for each donor and inducer combination. Haploid, hybrid, and defective kernels from the induction cross ears were manually sorted by experienced technical staff as described in section 2.3. A subset of the sorted kernels was placed into 24-well microtiter plates to track each kernel across all stages of this study.

**Table 1.** Counts of kernels for the three kernel classes from 12 induction populations.

| Population | Donor<sup>1</sup><br>♀ | Inducer<br>♂ | Kernel Count<sup>2</sup> |  |  | HIR<sub>v</sub> (%)<sup>3</sup> | HIR (%)<sup>4</sup> |
| --- | --- | --- | --- | --- | --- | --- | --- |
|  |  |  | Haploid | Hybrid | Defective |  |  |
| 1 | Viking 51 | IHO-1 | 32 | 154 | 36 | 9 | 8 |
| 2 | Viking 51 | IHO-2 | 35 | 146 | 10 | 13 | 13 |
| 3 | Viking 51 | IHO-3 | 34 | 153 | 0 | 10 | 10 |
| 4 | Viking 51 | IHO-4 | 35 | 157 | 41 | 11 | 11 |
| 5 | Viking 60 | IHO-2 | 31 | 154 | 24 | 10 | 10 |
| 6 | CCO-1 | IHO-1 | 23 | 158 | 16 | 6 | 4 |
| 7 | CCO-1 | IHO-3 | 13 | 174 | 7 | 3 | 3 |
| 8 | CCO-1 | IHO-4 | 28 | 151 | 11 | 8 | 8 |
| 9 | CCO-2 | IHO-1 | 43 | 144 | 0 | 8 | 8 |
| 10 | CCO-2 | IHO-2 | 38 | 149 | 12 | 12 | 9 |
| 11 | CCO-2 | IHO-3 | 36 | 151 | 12 | 9 | 8 |
| 12 | CCO-2 | IHO-4 | 45 | 138 | 12 | 12 | 12 |

<sup>1</sup>Viking 51 and Viking 60 are field corn donors; CCO-1 and CCO-2 are sweet corn donors
<sup>2</sup>Counts are based on gDNA scoring for haploid and hybrid kernels and visual scoring for defective kernels. Haploid and hybrid kernel counts reflect the number of kernels that were genotyped and are not based on the induction rates of each population
<sup>3</sup>Visual haploid induction rate (HIR<sub>v</sub>) is based on original visual sorting of each population
<sup>4</sup>Haploid induction rate (HIR) is based on original visual sorting of each population corrected by visual error rates (in Table 2) determined through gDNA scoring

### 2.2. Single-kernel near-infrared (skNIR) spectra collection and oil content predictions

NIR spectra were collected from kernels of 12 haploid induction populations (**Table 1**) using an automated skNIR sorter described by Graciano et al. (2025) and set to manual mode. Individual kernels were selected from the 24-well microtiter plate and manually dropped into an illuminated light tube. Fiber optic cables at the top and bottom of the light tube transmitted light to a CDI 256L-1.7T1 spectrophotometer, which features a 256-element InGaAs photodiode array spanning the spectral range from 906 to 1679 nm with a 1 nm step size (Control Development Inc., South Bend, IN). NIR reflectance values were trimmed to 940 to 1640 nm, and absorbance values were calculated as log(1/Reflectance). Oil content predictions for each kernel were calculated using partial least squares regression (PLSR) coefficients from a PLS model that was developed on the skNIR sorter using a well-established chemometric approach (Spielbauer et al. 2009; Gustin et al. 2013). As each kernel passed through the light tube, a two-way sorting stream guided it into one of two bins using a solenoid-based sorting system based on the kernel’s NIR-determined trait. Reference kernel oil content was determined as in Spielbauer et al. (2009) for 9 of the 12 induction populations using a PCT-20/20B NMR analyzer (Process Control Technology, Fort Collins, CO). Pure maize oil was used as the standard.

### 2.3. Ploidy verification

For each induction cross, kernels were visually scored by experienced technical staff as they were shelled from the ears into normal (typical grain-fill, embryo and scutellum development) or defective (atypical development; missing scutellum). Normal kernels were visually scored as haploid or hybrid using the *R1-nj* color marker. After skNIR data collection, 192 haploid and hybrid kernels per population were randomly selected for genotyping using gDNA markers. Haploid kernels were oversampled for genotyping relative to the HIR to ensure sufficient representation of the low-frequency class in subsequent analyses (**Table 1**). Kernels were germinated directly in their resident 24-well plate using hydrogel beads. Eighty-two of the selected kernels did not germinate and were not used in the study. Coleoptile and radicle tissue from each germinated kernel were used for DNA extraction. Ploidy was confirmed for each germinated kernel based on high-resolution melting curve analysis using primer pairs that discriminate between the wild type (donor) and mutated (inducer) alleles for the haploid induction gene *MATRILINEAL* (*Zm00001eb019170*) (Kelliher et al. 2017; Gilles et al. 2017; Liu et al. 2017). Of the 2,403 total kernels in the 12 induction populations, 393 were classified as haploid and 1,829 as hybrid by gDNA scoring, and 181 were classified as defective based on visual scoring (**Table 1**). The final counts of hybrid and haploid kernels reflected the genotyping sampling approach rather than the population HIR. Kernels from all parts of the ear and kernels with ambiguous/uncertain *R1-nj* coloration were included to ensure that the kernel set used for modeling was a representative sample of all kernels in the induction cross.

### 2.4. Metrics for model performance evaluation

Classification error rates were determined using two metrics, false discovery rate (FDR) and false negative rate (FNR), with haploid as the positive class and hybrid as the negative class for haploid/hybrid classification models and for evaluation of visual classification error. FDR is the proportion of predicted haploid kernels that were true hybrids, and FNR is the proportion of true haploid kernels incorrectly predicted as hybrids. True class labels were assigned by gDNA scoring. For all haploid/hybrid classification models described in sections 2.5 and 2.6, defective kernels were included only in the test set to simulate realistic sorting conditions but were excluded from FDR and FNR calculations. For the secondary defective kernel sorting step described in section 2.7, non-defective kernels (haploid and hybrid) were treated as the positive class and defective kernels as the negative class for metrics calculation.

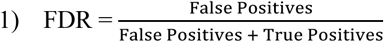

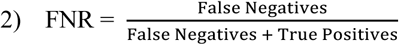

The HIR for each population was calculated from the visual classification of all kernels in the original population, corrected using the FDR and FNR of the visual classification (**Table 2**). The genotype-corrected haploid count (Ĥ) and subsequent HIR were obtained by solving:

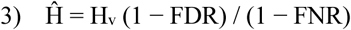

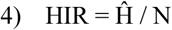

where Hv is the number of kernels visually classified as haploid and N is the total number of kernels visually classified as haploid or hybrid (excluding defective kernels) in the original population. HIRs calculated this way ranged from 3% to 13% across the 12 populations (**Table 1**).

**Table 2.** Classification error rates for manual haploid sorting using visual kernel color marker *R1-nj* compared to the gDNA scoring of haploid and hybrid kernels across 12 induction populations.

| Population | FDR<sup>1</sup> | FNR<sup>2</sup> |
| --- | --- | --- |
| Viking 51 × IHO-1 (1) | 0.11 | 0.03 |
| Viking 51 × IHO-2 (2) | 0.03 | 0.03 |
| Viking 51 × IHO-3 (3) | 0.03 | 0.00 |
| Viking 51 × IHO-4 (4) | 0.06 | 0.03 |
| Viking 60 × IHO-2 (5) | 0.11 | 0.00 |
| **Field Corn Mean** | **0.07** | **0.02** |
| CCO-1 × IHO-1 (6) | 0.26 | 0.00 |
| CCO-1 × IHO-3 (7) | 0.00 | 0.15 |
| CCO-1 × IHO-4 (8) | 0.11 | 0.14 |
| CCO-2 × IHO-1 (9) | 0.05 | 0.12 |
| CCO-2 × IHO-2 (10) | 0.19 | 0.00 |
| CCO-2 × IHO-3 (11) | 0.15 | 0.03 |
| CCO-2 × IHO-4 (12) | 0.09 | 0.09 |
| **Sweet Corn Mean** | **0.12** | **0.08** |
| **Overall Mean** | **0.10** | **0.05** |

<sup>1</sup>False discovery rate (FDR) is the proportion of predicted haploid kernels that were actually hybrids
<sup>2</sup>False negative rate (FNR) is the proportion of true haploid kernels that were incorrectly predicted as hybrids

### 2.5. Oil-based haploid classification

All classification analyses were performed on populations resampled to match the genotype-corrected HIR of each population to account for overrepresentation of haploid kernels in the pool selected for genotyping. Defective kernels were included in the resampled populations since they would be present under typical sorting conditions but were excluded from metric calculations. When resampling required a reduction in haploid kernel count to achieve the target HIR, defective kernels were downsampled at the same rate as haploid kernels to maintain realistic class proportions.

Two methods were evaluated for haploid sorting using predicted oil content (**Fig. 1**). First, the ‘lowest percentage’ method classified the X% of kernels with the lowest predicted oil content within each population as haploids, with X ranging from 4% to 20% in 2% increments. For each value of X, oil content values in the resampled population were sorted in ascending order, and the X^th^ percentile was used as the classification threshold. Second, the ‘dynamic threshold’ method classified kernels as haploids if their predicted oil content was below a threshold defined as the mean oil content of the resampled population minus a fixed offset value. Offset values of 0.8 to 2.4 were evaluated with a 0.2 step size for this approach. Neither approach attempted to classify defective kernels as a separate category; instead, defective kernels were sorted into the haploid or hybrid class based solely on their oil content. For each population and parameter value for the two methods, the population was resampled independently to its measured HIR in 20 separate iterations, and FDR and FNR were averaged across iterations to reduce variance in performance estimates. Optimal parameter values for the lowest percentage and dynamic threshold methods were selected by identifying the value that maximized the number of populations meeting both the FDR ≤ 0.25 and FNR ≤ 0.50 operational benchmarks, with mean FDR used as a tiebreaker. These benchmarks were established based on the practical experience of the Iowa State University Doubled Haploid Facility and reflect the asymmetric resource costs of classification errors in a DH production pipeline. Hybrid kernels advancing past the sorting stage (FDR) require chromosome doubling, greenhouse, and genotyping resources without contributing usable DH lines (FDR), whereas haploid kernels lost to the hybrid pool (FNR) reduce yield but can be compensated for by scaling induction population size.

**Fig. 1.**
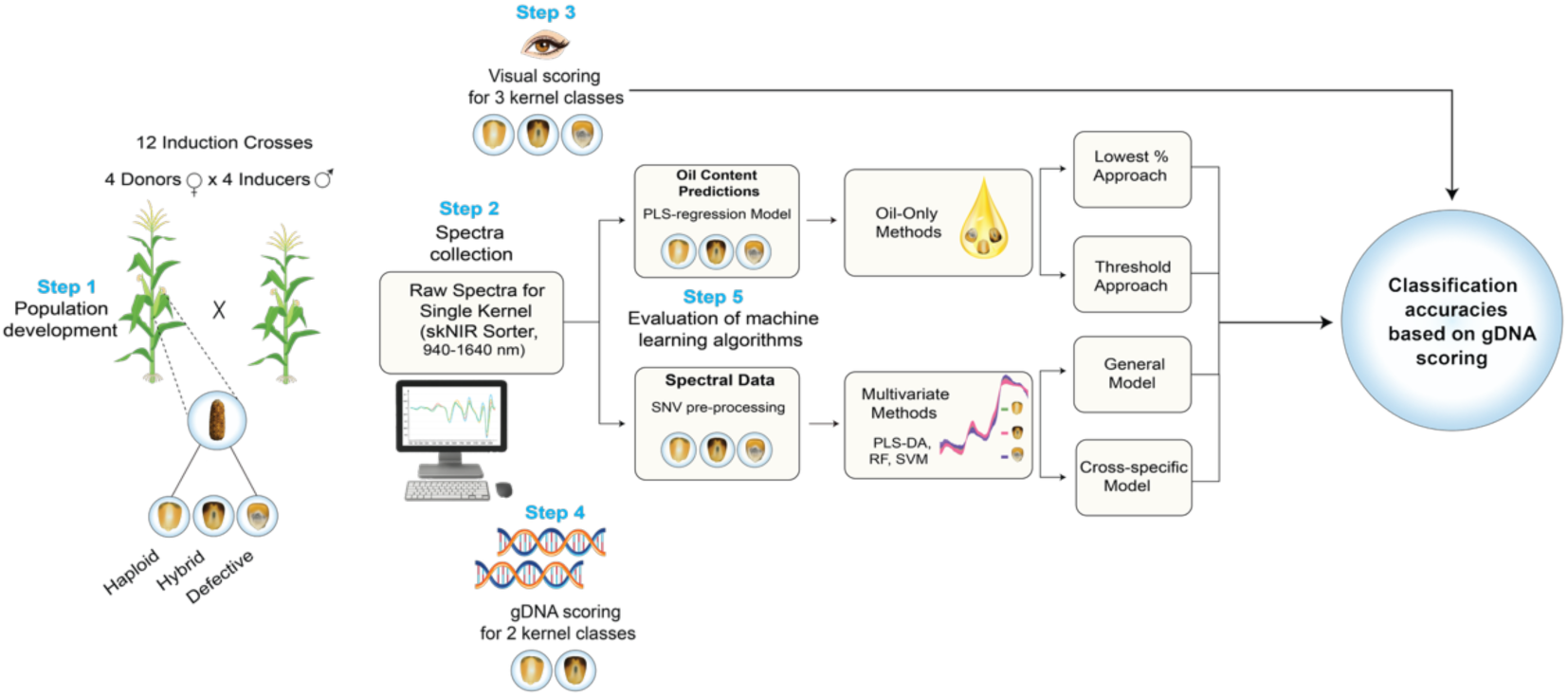
Schematic workflow from spectral data collection to evaluation of haploid sorting approaches using single-kernel near-infrared reflectance spectroscopy sorter.

To establish a performance ceiling for oil-based sorting methods, a ‘population-specific optimal threshold’ was determined for each population by evaluating 500 random candidate oil threshold values spanning the full range of predicted oil content. At each threshold value, the population was resampled to its target HIR over five independent iterations, and mean FDR and FNR values were calculated. Defective kernels were included in the resampled population for threshold derivation but excluded from FDR and FNR calculation. The threshold minimizing FNR subject to FDR ≤ 0.25 and FNR ≤ 0.50 for the given population was selected as the ‘per-population optimal threshold’. If no threshold satisfied both criteria, the threshold minimizing the combined metric (FDR + FNR) was selected.

### 2.6. Multivariate methods for haploid identification

Multivariate classification models were developed using NIR spectra from kernels of each induction cross (**Fig. 1**). Spectra were normalized to a mean of 0 and a standard deviation of 1 using the standard normal variate pretreatment implemented in the R package *waves* (v0.2.5; Hershberger et al. 2021). Three supervised machine learning multivariate methods were evaluated for binary classification: partial least squares discriminant analysis (PLS-DA), random forest (RF), and support vector machine (SVM) using the R package *caret* (v7.0-1; Kuhn 2008) with the “pls”, “rf”, and “svmRadial” methods, respectively. Model training employed 5-fold cross-validation to optimize performance, and Cohen’s Kappa was used as the evaluation metric. Within each iteration, a fixed random seed was applied to ensure reproducibility, and the same training and test sets were used to compare model performance of the three multivariate methods.

PLS-DA models were fit using 12 optimal latent components after tuning (ncomp = 12). SVM models used a radial basis function kernel to capture non-linear relationships in the high-dimensional spectral data with cost and sigma parameters tuned over a grid of cost = {1, 10, 100} and sigma = {0.0001, 0.001, 0.01}. RF models were constructed with 200 trees with the mtry parameter tuned over a grid of {10, 50, 100, √*p*} where *p* is the number of spectral features, using 50 preliminary trees per candidate value to select the optimal mtry. The Clemson Palmetto Cluster was used to perform the analyses in R v4.4.0 (Antao et al. 2024; R Core Team 2024)

As described in section 2.5, test sets were resampled to match the measured HIR of each population and defective kernels were included in the resampled test sets. Defective kernels were excluded from model training in all analyses to maximize the distinction between haploid and hybrid classes but were included in the test set to evaluate classification behavior under realistic sorting conditions. FDR and FNR were calculated on haploid and hybrid kernels only, excluding defective kernels from metric calculation.

A general model was trained on combined spectral data from all 12 induction populations using hold-one-population-out cross-validation (HOPO-CV). In each HOPO-CV round, all haploid and hybrid kernels from the 11 non-held-out populations were used for training. The held-out test population was resampled to its own measured HIR. The downsampled defective kernels from the held-out population were included in the test set. The full HOPO-CV procedure was repeated 20 times with independent resampling of the test set in each replication, and FDR and FNR were averaged across replications to produce stable performance estimates.

Population-specific models were also developed for each of the 12 induction populations to provide a performance ceiling for comparison. For each population, haploid and hybrid kernels were resampled to the population’s measured HIR and then split into 70% training and 30% test sets by stratified random sampling. Defective kernels were downsampled at the same rate as haploids when haploid downsampling was required, and 30% of the downsampled defectives were included in the test set. This procedure was repeated for 100 independent iterations with independent resampling within each iteration, and FDR and FNR were averaged across iterations.

For reference, populations achieving both operational benchmarks of FDR ≤ 0.25 and FNR ≤ 0.50 under the population-specific PLS-DA model are indicated in the results. This threshold represents the maximum classification accuracy achievable from skNIR spectral data for each population and provides a useful reference point for evaluating general model performance across diverse donor and inducer combinations.

### 2.7. Defective kernel sorting

A secondary sorting model was developed to remove defective kernels from the predicted haploid pool following the initial haploid sorting step. Operationally, this two-step approach first sorts kernels into a predicted haploid pool using either the best performing oil-based approach or the selected general multivariate model and then passes the smaller predicted haploid pool through a defective kernel cleaning step to remove any defective kernels that were incorrectly classified as haploid in the first sorting step. The defective kernel sorting was evaluated using a binary classification framework in which all haploid and hybrid kernels were relabeled as non-defective and defective kernels retained their class label.

Ten populations containing defective kernels (populations 3 and 9 were excluded as they contained no defective kernels) were evaluated using the HOPO-CV described previously in the general haploid/hybrid classification model. In each HOPO-CV fold, all haploid and hybrid kernels from 11 non-held-out populations were included as the non-defective class for training, while all defective kernels from those populations were included as the defective class. The test set for the held-out population was composed to reflect the expected composition of the predicted haploid pool following the initial sorting step. All haploid and defective kernels from the held-out population were included in the test set. Hybrid kernels were sampled to produce a dataset composition of 75% haploid and 25% hybrid kernels, reflecting the benchmark FDR of 0.25 or lower established for the initial sorting step, where up to 25% of kernels predicted as haploids could actually be hybrids. The same three multivariate classification methods evaluated for haploid sorting were applied to the defective kernel sorting task using the same model parameters. The full HOPO-CV procedure was repeated 20 times with independent resampling of hybrid kernels in each replication, and FDR and FNR were averaged across replications to produce stable performance estimates.

### 2.8. Statistical analysis

Significant differences in the predicted oil content (% fresh weight (f.w.)) among the three kernel classes were evaluated independently within each induction population including all the kernels used in the study (**Table 1**). Data normality for each kernel class was checked with the Shapiro-Wilk test using the R package *rstatix* (v0.7.2; Kassambara 2023). A non-parametric test (Kruskal-Wallis) was performed for each induction cross because the predicted oil content for each kernel class did not meet the assumptions of normal distribution. The Kruskal-Wallis test determines significant differences between the medians of more than two independent groups by ranking all the data points together in ascending order and comparing the average ranks across groups (Chan and Walmsley 1997). After significant differences were detected for each population (α = 0.05), post-hoc comparisons were conducted using Dunn’s test with Bonferroni correction to adjust for the multiple comparisons using packages *FSA* (v0.9.6; Ogle et al. 2025) and *multcompView* (v0.1-11; Graves et al. 2024). The packages *ggplot2* (v4.0.3; Wickham 2016), *cowplot* (v1.2.0; Wilke 2025), *ggrepel* (v0.9.6; Slowikowski 2026), and *ggtext* (v0.1.2; Wilke and Wiernik 2022) were used to create data visualizations.

The delta oil (Δ oil) for each population was calculated as a difference in mean predicted oil content between hybrid and haploid kernel classes. To evaluate genetic effects of parent backgrounds on Δ oil and HIR, two-way analysis of variance (ANOVA) was performed with donor and haploid inducer as the two fixed factors. The design was unreplicated because each of the 12 donor × inducer population combinations was represented by a single observation; therefore, the interaction term could not be independently estimated. Type III sum of squares were used for all tests to account for the unbalanced structure of the dataset (12 of 16 possible populations represented), implemented in R using the *car* package (v3.1-5; Fox and Weisberg 2019). The proportion of total variance attributable to each factor was quantified using eta-squared (η²), calculated as the ratio of each factor’s sum of squares to the total model sum of squares (inducer SS + donor SS + residual SS). To rank individual donors and inducers within each metric, least squares means (LS means) were estimated from each model using the *emmeans* package (v2.0.3; Lenth and Piaskowski 2026). LS means account for the unequal representation of donors and inducers across combinations and provide unbiased marginal estimates of performance for each level. Pairwise comparisons among donors and among inducers were conducted using Bonferroni correction to adjust for the multiple comparisons.

The skNIR sorter’s PLS model performance for oil content predictions was evaluated by calculating the coefficient of determination (R^2^) and root mean squared error (RMSE) compared to the NMR-measured values. Pearson’s correlation coefficients (*r*) and *p*-values were calculated to determine the degree of relationship between two continuous variables using the built-in *stats* package in R.

## 3. Results

### 3.1. Variation in the NIR spectra and predicted oil content for kernel classes in 12 induction populations

#### 3.1.1. Differences in the NIR spectra for the three kernel classes

NIR-based discrimination of haploid from hybrid kernels requires chemical and/or structural differences between the classes, resulting in predictable variation among NIR spectra. Only slight differences were observed for the mean NIR spectra of haploid and hybrid kernels when all induction crosses were combined, while defective kernels revealed a distinct spectral profile (**Fig. 2a)**. Within individual induction crosses, however, haploid and hybrid spectral profiles exhibited greater divergence, with the most pronounced class separation occurring near the lower and upper limits of the measured wavelength range (940 nm and 1640 nm, respectively) for most populations (**Fig. 2b** and **Supplementary Fig. 1**).

**Fig. 2.**
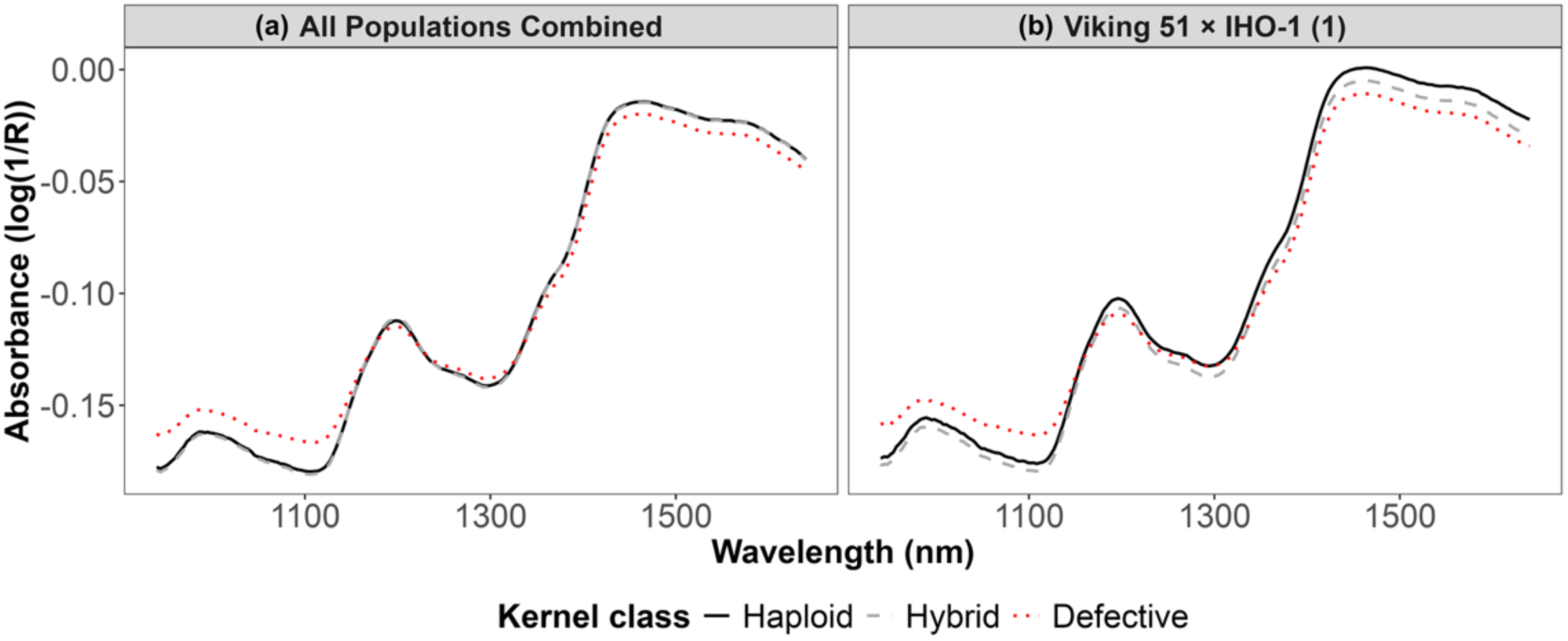
Raw near-infrared (NIR) spectra of kernels resulting from 12 induction populations in field and sweet corn backgrounds, **(a)** Mean absorbance spectra of haploid, hybrid and defective kernels with all induction crosses combined, **(b)** Mean absorbance spectra of haploid, hybrid, and defective kernels from the induction population 1 (Viking 51 × IHO-1). Absorbance is reported as log(1/Reflectance) for the NIR wavelength range of 940 to 1640 nm using single-kernel NIR spectroscopy sorter.

#### 3.1.2. skNIR-predicted oil content variation for the three kernel classes

The oil content of kernels from 12 induction crosses was estimated using a PLS model implemented on the skNIR system, which showed expected predictive ability based on published PLS model performance and the NMR-measured oil content across the three kernel classes with an RMSE of 0.65% for haploid kernels, 0.96% for hybrid kernels, and 1.77% for defective kernel predictions (**Supplementary Fig. 2**).

Across all donor and inducer combinations, hybrid kernels exhibited significantly higher predicted oil content than haploid or defective kernels (**Fig. 3**). Defective kernels showed the lowest oil content, but distribution was not significantly different from haploid kernels within each induction cross. The sweet corn donor CCO-1 produced the highest haploid and hybrid oil content distributions across all of its induction cross combinations, followed by the sweet corn donor CCO-2, but with more overlapping distributions for the two kernel classes. Compared with the sweet corn donors, field corn donors Viking 51 and Viking 60 showed consistently lower oil content distributions for the kernel classes.

**Fig. 3.**
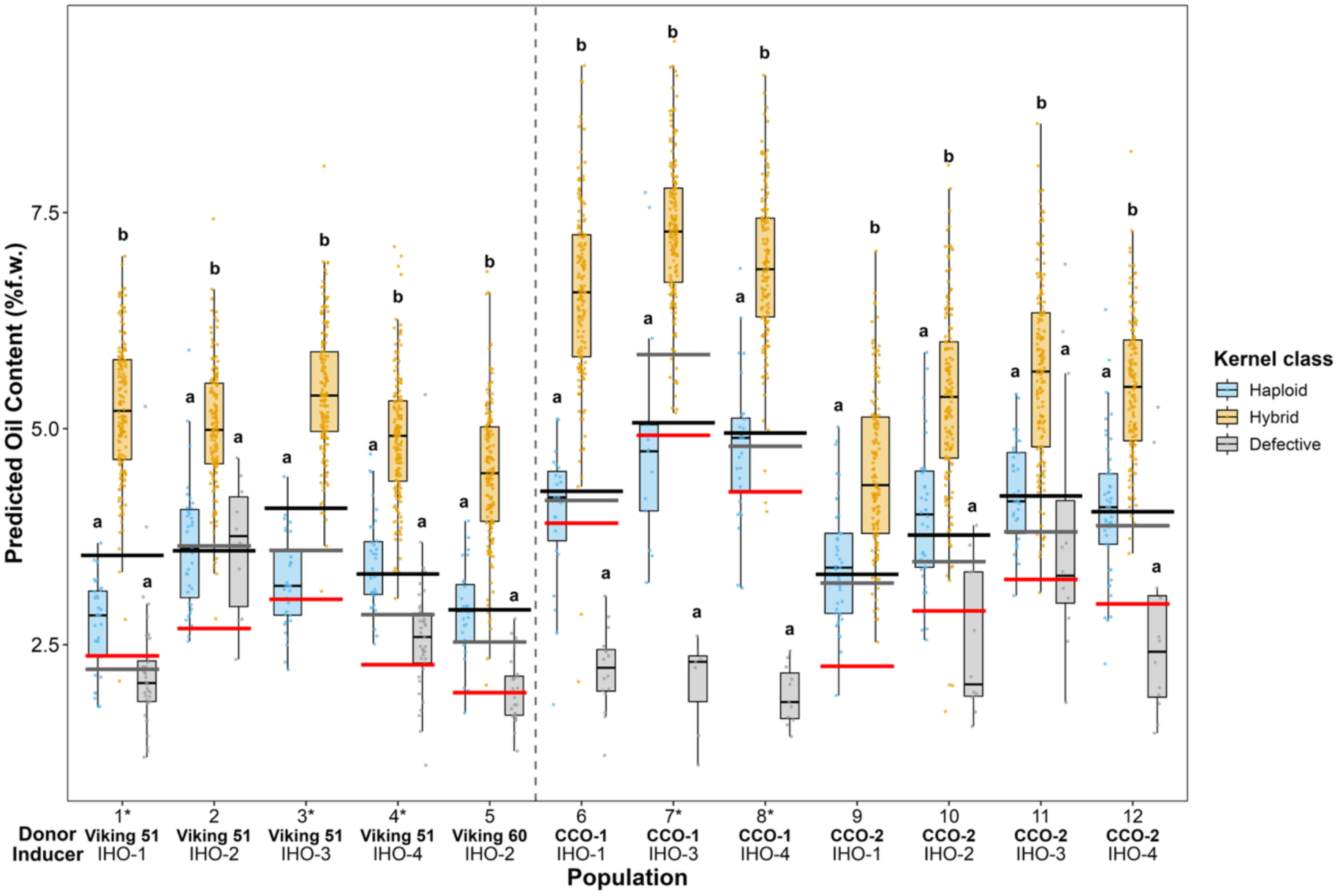
Predicted oil content distributions for the three kernel classes across 12 induction populations. Each colored boxplot represents a kernel class within an induction cross and contains jittered points for individual kernels used in the study. Among the three boxplots for each population, distributions with the same letter are not significantly different based on Dunn’s test with Bonferroni correction pairwise comparisons. The black dashed vertical line separates field corn populations (left; populations 1-5) from sweet corn populations (right; populations 6-12). Black horizontal lines indicate the ‘per-population optimal’ classification threshold for each induction cross based on resampling-based optimization procedure. The gray horizontal lines indicate the classification threshold for the ‘lowest percentage’ method (14^th^ percentile; selected threshold from Table 3) for each induction population. Red horizontal lines indicate the classification threshold for the ‘dynamic threshold’ method (offset = 2.0; selected offset from Table 3) for each induction population. Asterisks (*) indicate populations that met the FDR ≤ 0.25 and FNR ≤ 0.50 operational benchmarks under the per-population optimal threshold.

**Table 3.**
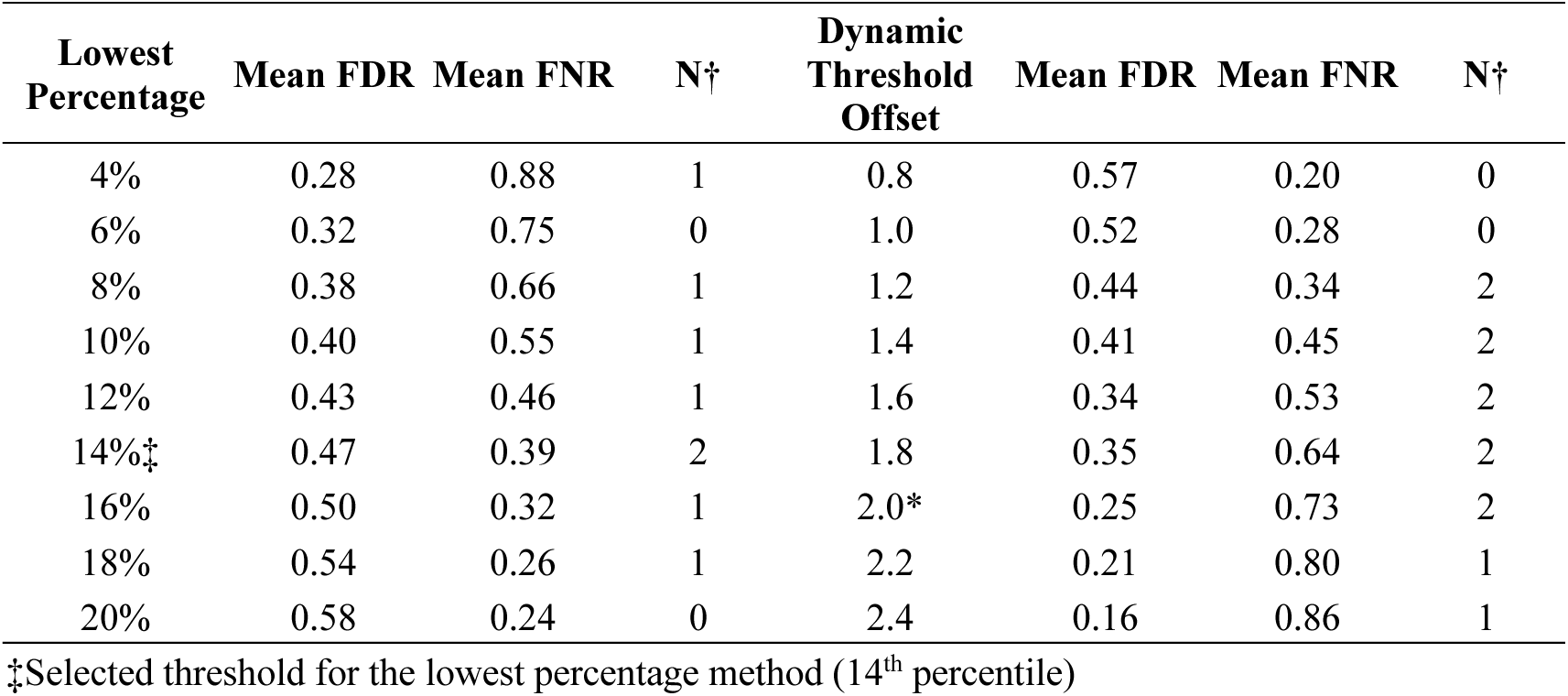
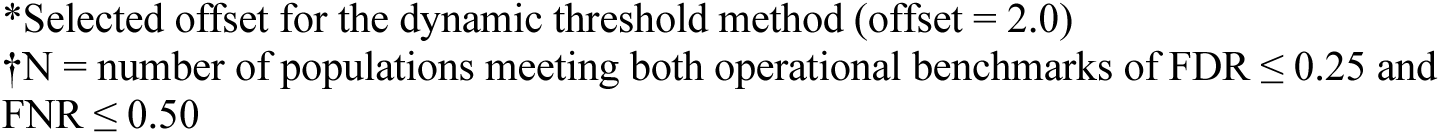
Mean false discovery rate (FDR) and false negative rate (FNR) for 12 induction populations across a range of the lowest percentage and dynamic threshold offset parameter values for oil-based methods.

| Lowest Percentage | Mean FDR | Mean FNR | N† | Dynamic Threshold Offset | Mean FDR | Mean FNR | N† |
| --- | --- | --- | --- | --- | --- | --- | --- |
| 4% | 0.28 | 0.88 | 1 | 0.8 | 0.57 | 0.20 | 0 |
| 6% | 0.32 | 0.75 | 0 | 1.0 | 0.52 | 0.28 | 0 |
| 8% | 0.38 | 0.66 | 1 | 1.2 | 0.44 | 0.34 | 2 |
| 10% | 0.40 | 0.55 | 1 | 1.4 | 0.41 | 0.45 | 2 |
| 12% | 0.43 | 0.46 | 1 | 1.6 | 0.34 | 0.53 | 2 |
| 14%‡ | 0.47 | 0.39 | 2 | 1.8 | 0.35 | 0.64 | 2 |
| 16% | 0.50 | 0.32 | 1 | 2.0* | 0.25 | 0.73 | 2 |
| 18% | 0.54 | 0.26 | 1 | 2.2 | 0.21 | 0.80 | 1 |
| 20% | 0.58 | 0.24 | 0 | 2.4 | 0.16 | 0.86 | 1 |

‡Selected threshold for the lowest percentage method (14<sup>th</sup> percentile)
\*Selected offset for the dynamic threshold method (offset = 2.0) $\dagger N$ = number of populations meeting both operational benchmarks of $FDR \leq 0.25$ and $FNR \leq 0.50$

The relative difference in predicted oil content between the hybrid and haploid classes within each induction population (Δ oil) was used to evaluate the class discriminating power of each inducer (**Fig. 4**). Overall, IHO-1 and IHO-3 generated the largest Δ oil, while IHO-2 populations had the smallest (**Fig. 4**). Mean Δ oil across all 12 populations was 1.77%, ranging from 2.54% in CCO-1 × IHO-1 (6) to 1.07% in CCO-2 × IHO-1 (9). A two-way ANOVA was used to partition the contributions of donor and inducer to Δ oil variation (**Supplementary Tables 1 and 2**). Donor was the dominant source of variation, accounting for 60.3% of total variance (η² = 0.603, *p* = 0.079), compared to 15.6% for inducer (η² = 0.156, *p* = 0.438) (**Supplementary Table 1**). Among donors, CCO-1 had the largest LS means for Δ oil (2.24%) and CCO-2 the smallest (1.28%), consistent with the class separation observed in **Fig. 3**. Among inducers, IHO-1 generated the largest LS means for Δ oil (2.04%) and IHO-2 the smallest (1.58%), though the range was narrower than among donors (**Supplementary Table 2**). Bonferroni-adjusted pairwise comparisons showed overlapping confidence intervals for all donors and all inducers, with no pairs showing a statistically significant difference (*p* > 0.05). The lack of statistical significance is attributable to limited statistical power, with only 5 error degrees of freedom in this unreplicated design.

**Fig. 4.**
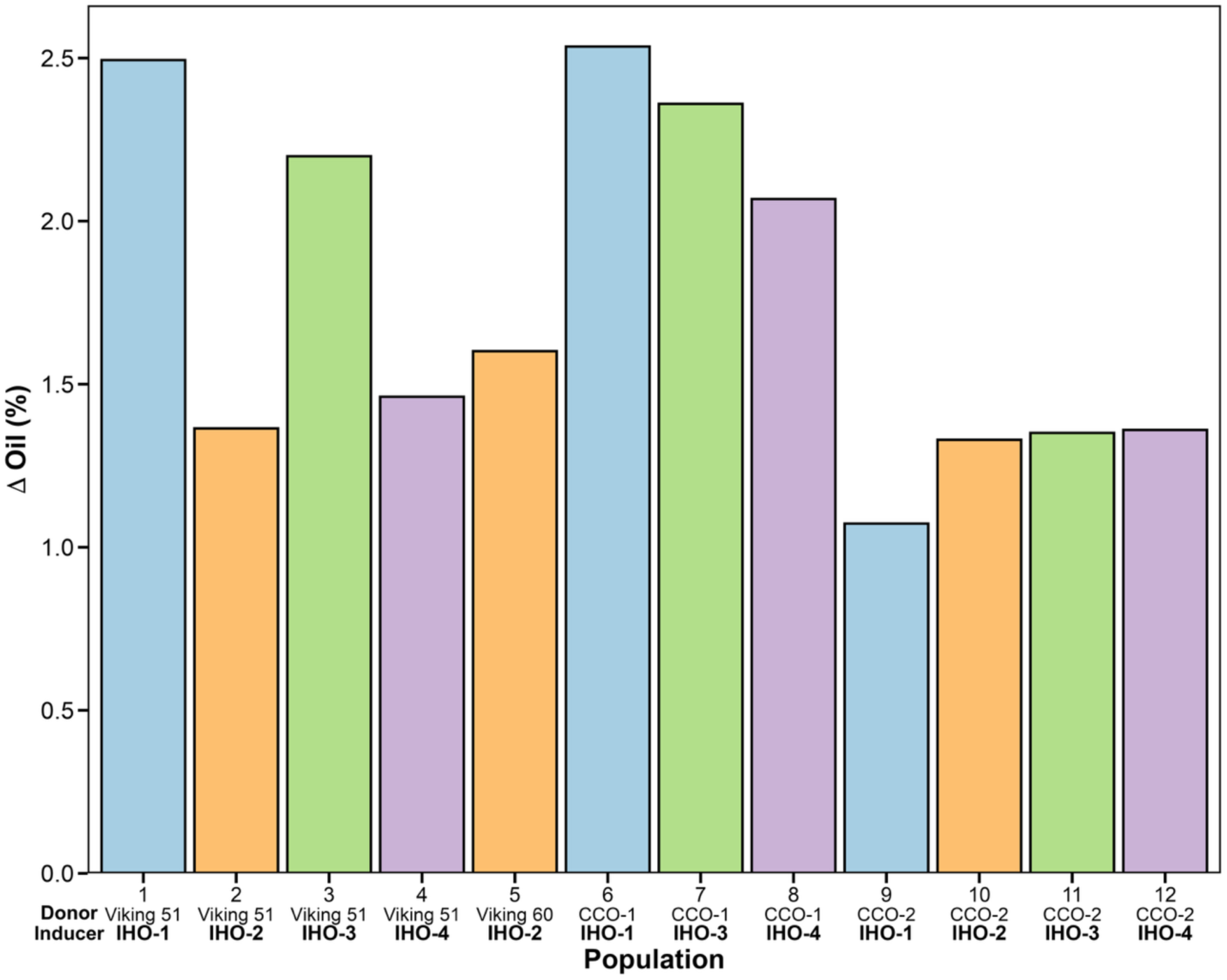
Effects of four high-oil haploid inducers (IHO-1, IHO-2, IHO-3, and IHO-4) on Δ oil (mean hybrid oil content – mean haploid oil content) of each induction population. Viking 51 and Viking 60 are field corn donors and CCO-1 and CCO-2 are sweet corn donors. Bars are color-coded by inducer, with the same color representing the same inducer across different crosses.

HIR was analyzed using the same two-way ANOVA framework (**Supplementary Tables 1 and 2**). Donor accounted for 54.0% of total HIR variance (η² = 0.540, *p* = 0.026) and inducer accounted for 34.2% (η² = 0.342, *p* = 0.062) (**Supplementary Table 1**). Among donors, CCO-1 had the lowest LS means for HIR (5.38%) and Viking 51 had the highest (10.50%), with Viking 51 showing significantly higher HIR than CCO-1 in Bonferroni-adjusted pairwise comparisons (*p* = 0.033) (**Supplementary Table 2**). Among inducers, IHO-1 and IHO-3 had the lowest LS means for HIR (6.79% and 7.13%, respectively) while IHO-4 had the highest (10.46%), though no pairwise comparisons among inducers were significantly different (*p* > 0.05). Moreover, IHO-1 and IHO-3 (the inducers generating the largest Δ oil) also showed the lowest HIR, while IHO-2 (the inducer generating the smallest Δ oil) showed the second highest HIR. This inverse relationship is supported by a significant negative correlation (*r* = −0.62, *p* = 0.032) between Δ oil generation and HIR among the IHO lines.

### 3.2. Visual marker accuracy for haploid discrimination

Manual haploid sorting based on the *R1-nj* kernel color marker was evaluated against genetic marker-based scoring to assess visual sorting accuracy. Overall, the average FDR and FNR for visual classification were 0.10 and 0.05, respectively (**Table 2**). Two field corn induction crosses, Viking 51 × IHO-2 (2) and Viking 51 × IHO-3 (3), had near-perfect visual separation of haploid kernels. However, visual sorting error rates were consistently higher across sweet corn populations than field corn, with mean error rates approximately double that of the field corn induction populations. Two populations involving the sweet corn donor CCO-1, CCO-1 × IHO-1 (6) and CCO-1 × IHO-4 (8), had the highest visual sorting error rates (FDR + FNR = 0.26 and 0.25, respectively), indicating relatively stronger anthocyanin suppression in CCO-1.

### 3.3. NIR-based haploid sorting using predicted oil content

Predicted oil content derived from skNIR spectra was evaluated as the basis for haploid kernel sorting across 12 induction populations representing 4 donor backgrounds (Viking 51, Viking 60, CCO-1 and CCO-2) crossed with four experimental inducers (IHO-1 through IHO-4). Two general sorting methods were developed and evaluated: a lowest percentage method that classifies the bottom X% of kernels within each population as haploid based on predicted oil content, and a dynamic threshold method that classifies kernels as haploid if their predicted oil content falls below the population mean minus a fixed offset. Both methods were designed for deployment on the skNIR sorter without requiring population-specific calibration or prior knowledge of kernel class composition.

#### 3.3.1 Per-population optimal threshold analysis

To establish a performance ceiling for oil content-based sorting, a per-population optimal threshold analysis was conducted by evaluating 500 random classification thresholds across the full range of observed oil content values for each population and identifying the threshold that minimized FNR subject to the operational benchmarks FDR ≤ 0.25 and FNR ≤ 0.50.

Per-population optimal analysis identified five populations for which FDR ≤ 0.25 was achievable within the FNR ≤ 0.50 constraint: Viking 51 × IHO-1 (1), Viking 51 × IHO-3 (3), Viking 51 × IHO-4 (4), CCO-1 × IHO-3 (7) and CCO-1 × IHO-4 (8) (**Supplementary Table 3**). These five populations meeting the operational benchmarks were used for general threshold parameter selection. The remaining seven populations could not simultaneously meet both operational benchmarks under any oil content threshold. Among all 12 populations, both mean FDR and FNR at the per-population optimal threshold were 0.33.

The five populations achieving both operational benchmarks included three of the four Viking 51 donor populations and two of the three CCO-1 populations, while all CCO-2 populations and the Viking 60 population failed to meet both benchmarks under any oil content threshold. The failure of CCO-2 and Viking 60 populations is consistent with their lower Δ oil values relative to the Viking 51 and CCO-1 populations (**Fig. 4**). The Viking 51 and CCO-1 populations crossed with the inducers that drove the largest Δ oil, IHO-1 and IHO-3, all met the FDR and FNR benchmarks with the exception of CCO-1 × IHO-1 (6), suggesting these inducers show particular promise for producing populations amenable to oil-based haploid sorting. CCO-1 × IHO-1 (6) had the second lowest HIR (4%) of all 12 populations, which likely contributed to its operationally unacceptable FDR of 0.33 (**Supplementary Table 3**) despite having the largest Δ oil (2.54%) of all 12 populations (**Fig. 4**). CCO-1 × IHO-3 (7) had the lowest HIR (3%) but did meet the benchmark showing that low HIR does not preclude populations from accurate population-specific sorting.

#### 3.3.2 General threshold selection and evaluation of two oil-based methods

General threshold parameters for the lowest percentage and dynamic threshold methods were selected by identifying the parameter values that maximized the number of populations meeting both FDR ≤ 0.25 and FNR ≤ 0.50, with mean FDR used as a tiebreaker (**Table 3**).

For the lowest percentage method, sweeping the classification percentile from 4% to 20% in 2% increments revealed a consistent tradeoff between FDR and FNR across all 12 populations (**Table 3**). The 14^th^ percentile was selected as the optimal threshold, with two of 12 populations meeting both operational benchmarks (Viking 51 × IHO-1 (1) and Viking 51 × IHO-3 (3)) and a mean FDR of 0.47 and mean FNR of 0.39 across all 12 populations (**Supplementary Table 3**). For the dynamic threshold method, sweeping the offset from 0.8 to 2.4 in increments of 0.2 similarly revealed an FDR-FNR tradeoff (**Table 3**). An offset of 2.0 was selected as optimal, with two of 12 populations meeting both operational benchmarks (Viking 51 × IHO-3 (3) and CCO-1 × IHO-3 (7)) and a mean FDR of 0.25 and mean FNR of 0.73 across all 12 populations (**Supplementary Table 3**).

The selected offset for the dynamic threshold method tended to set the threshold closer to the defective class than either the per-population optimal or the lowest percentage threshold, resulting in a relatively cleaner haploid pool (lower FDR) at the expense of higher loss of haploids into the predicted hybrid pool (higher FNR) (**Fig. 3**). The dynamic threshold method had lower mean FDR than the lowest percentage method (0.25 vs. 0.47, respectively) while substantially higher mean FNR (0.73 vs. 0.39, respectively), reflecting the inherent tradeoff between the two error types (**Supplementary Table 3**). Reducing the dynamic threshold offset from 2.0 to 1.6 produced a more balanced FDR and FNR but did not improve the number of populations meeting both FDR ≤ 0.25 and FNR ≤ 0.50 (**Table 3**).

Viking 51 × IHO-3 (3) was the only population meeting both operational benchmarks under both general methods (**Supplementary Table 3**). Viking 51 × IHO-1 (1) narrowly missed the FNR ≤ 0.50 benchmark under the dynamic threshold method (FNR = 0.53), and CCO-1 × IHO-3 (7) showed elevated FDR under the lowest percentage method (FDR = 0.79), reflecting the difficulty of applying a fixed percentage threshold to a population with a very low HIR of 3% where defective kernels represent a disproportionate share of the lowest oil content kernels. IHO-1 and IHO-3 were the inducers in most of the populations meeting both benchmarks under at least one general method, consistent with these inducers generating the largest Δ oil across donor backgrounds.

The CCO-2 donor populations consistently failed to meet both operational benchmarks under any oil-based method, with mean FDR and FNR well above the operational thresholds for the two methods and per-population optimal threshold analysis. The CCO-2 populations with IHO-1 and IHO-3 inducers were no more accurate than the IHO-2 and IHO-4 inducers showing that inducers that drive larger Δ oil in general did not improve sorting accuracy in the CCO-2 donor background.

### 3.4. Multivariate spectral classification for haploid sorting

Haploid discrimination accuracy may be improved by exploiting the full skNIR spectral profile beyond what is achievable using predicted oil content alone. Three supervised multivariate classification methods were evaluated for haploid kernel discrimination across the 12 induction populations: PLS-DA, SVM, and RF. Models were evaluated under two configurations: population-specific models trained and tested within each induction population to establish a performance ceiling, and a general model trained across all 12 populations and evaluated using HOPO-CV as the deployable sorting approach.

#### 3.4.1. Population-specific model performance

Population-specific models were evaluated first to establish the upper bound of classification accuracy achievable with multivariate NIR spectral methods. Because population-specific models require prior genotyping of each target population, they are not ideal for operational deployment but provide a reference for the best achievable classification performance using multivariate spectral data.

Overall, PLS-DA had the lowest mean FNR among the population-specific models, while FDR remained comparable across the three multivariate methods (0.22 to 0.25) (**Table 4**). Based on the operational benchmarks of FDR ≤ 0.25 and FNR ≤ 0.50, population-specific PLS-DA achieved both benchmarks in five of the 12 induction populations (Viking 51 × IHO-1 (1), Viking 51 × IHO-2 (2), Viking 51 × IHO-3 (3), Viking 51 × IHO-4 (4), and Viking 60 × IHO-2 (5)) (**Table 4**) and included all of the Viking donor populations. Similarly, the population-specific SVM also had four Viking 51 donor populations meeting operational benchmarks, compared with five populations for the PLS-DA. CCO-1 × IHO-4 (8) narrowly missed the FDR ≤ 0.25 benchmark with FDR = 0.28 and 0.26 under population-specific PLS-DA and SVM, respectively. Viking 51 × IHO-2 (2) and Viking 51 × IHO-3 (3) achieved near-perfect classification under population-specific PLS-DA with FDR = 0.08, FNR = 0.05 and FDR = 0.05, FNR = 0.01, respectively. The remaining seven populations showed substantially higher classification error rates under population-specific PLS-DA, with all CCO-2 donor populations showing mean FDR > 0.30 and mean FNR > 0.50, suggesting that this donor background generates insufficient spectral differentiation between haploid and hybrid kernels to support reliable classification by any NIR-based method evaluated here.

**Table 4.** Population-specific model classification error rates for three multivariate methods across 12 induction populations.

| Population | PLS-DA<sup>‡</sup> |  | SVM<sup>‡</sup> |  | RF<sup>‡</sup> |  |
| --- | --- | --- | --- | --- | --- | --- |
|  | FDR<sup>1</sup> | FNR<sup>2</sup> | FDR<sup>1</sup> | FNR<sup>2</sup> | FDR<sup>1</sup> | FNR<sup>2</sup> |
| Viking 51 × IHO-1 (1)* | **0.22** | **0.25** | **0.21** | **0.35** | 0.31 | 0.74 |
| Viking 51 × IHO-2 (2)* | **0.08** | **0.05** | **0.11** | **0.15** | 0.23 | 0.64 |
| Viking 51 × IHO-3 (3)* | **0.05** | **0.01** | **0.01** | **0.12** | **0.09** | **0.37** |
| Viking 51 × IHO-4 (4)* | **0.10** | **0.09** | **0.02** | **0.18** | 0.08 | 0.74 |
| Viking 60 × IHO-2 (5)* | **0.17** | **0.20** | 0.27 | 0.53 | 0.28 | 0.95 |
|  | FDR<sup>1</sup> | FNR<sup>2</sup> | FDR<sup>1</sup> | FNR<sup>2</sup> | FDR<sup>1</sup> | FNR<sup>2</sup> |
| CCO-1 × IHO-1 (6) | 0.21 | 0.66 | 0.27 | 0.80 | 0.18 | 0.81 |
| CCO-1 × IHO-3 (7) | 0.00 | 0.79 | 0.06 | 0.62 | 0.05 | 0.66 |
| CCO-1 × IHO-4 (8) | 0.28 | 0.49 | 0.26 | 0.46 | 0.26 | 0.41 |
| CCO-2 × IHO-1 (9) | 0.51 | 0.64 | 0.41 | 0.85 | 0.12 | 0.97 |
| CCO-2 × IHO-2 (10) | 0.54 | 0.75 | 0.49 | 0.82 | 0.30 | 0.95 |
| CCO-2 × IHO-3 (11) | 0.53 | 0.71 | 0.54 | 0.74 | 0.38 | 0.80 |
| CCO-2 × IHO-4 (12) | 0.36 | 0.58 | 0.34 | 0.58 | 0.40 | 0.74 |
| **Mean** | **0.25** | **0.43** | **0.25** | **0.52** | **0.22** | **0.73** |

<sup>‡</sup>Three multivariate methods: partial least squares discriminant analysis (PLS-DA), support vector machine (SVM), and random forest (RF)
<sup>1</sup>False discovery rate (FDR) is the proportion of predicted haploid kernels that were actually hybrids
<sup>2</sup>False negative rate (FNR) is the proportion of true haploid kernels that were incorrectly predicted as hybrids
\*Population met $FDR \leq 0.25$ and $FNR \leq 0.50$ operational benchmarks under population-specific PLS-DA model, providing a reference for the best achievable classification performance using multivariate spectral data
Bolded values indicate populations that passed the $FDR \leq 0.25$ and $FNR \leq 0.50$ operational benchmarks under the given multivariate population-specific model. Population-specific models were trained and evaluated within each induction population (70% training (haploids and hybrids) and 30% test set (haploids, hybrids and defectives))

#### 3.4.2. General model performance

The general SVM model achieved the best overall performance among the three general model methods, with mean FDR = 0.32 and mean FNR = 0.38 across all 12 populations (**Table 5**). Both general model PLS-DA and RF showed comparable results with mean FDR of 0.38 and 0.35 and mean FNR of 0.45 and 0.44, respectively. The general SVM met both operational benchmarks in four of the 12 populations (Viking 51 × IHO-1 (1), Viking 51 × IHO-2 (2), Viking 51 × IHO-3 (3), and CCO-1 × IHO-3 (7)), three of which were also identified as sortable under population-specific PLS-DA (**Table 4**). Particularly, the overall mean FDR and FNR were relatively similar between the population-specific PLS-DA (mean FDR = 0.25, mean FNR = 0.43) and the general SVM (mean FDR = 0.32, mean FNR = 0.38), suggesting that the general model achieves comparable overall classification accuracy to the population-specific ceiling despite requiring no prior genotyping. Like the other classification approaches, the FDR and FNR varied substantially across populations under the general SVM suggesting that the ability of the general model to recover haploids is also highly population-dependent and likely influenced by donor background and HIR.

**Table 5.** General model classification error rates for three multivariate methods across 12 induction populations.

| Population | PLS-DA<sup>‡</sup> |  | SVM<sup>‡</sup> |  | RF<sup>‡</sup> |  |
| --- | --- | --- | --- | --- | --- | --- |
|  | FDR<sup>1</sup> | FNR<sup>2</sup> | FDR<sup>1</sup> | FNR<sup>2</sup> | FDR<sup>1</sup> | FNR<sup>2</sup> |
| Viking 51 × IHO-1 (1)* | 0.33 | 0.06 | **0.15** | **0.11** | **0.25** | **0.20** |
| Viking 51 × IHO-2 (2)* | 0.29 | 0.55 | **0.18** | **0.39** | 0.18 | 0.58 |
| Viking 51 × IHO-3 (3)* | **0.00** | **0.44** | **0.02** | **0.44** | **0.00** | **0.43** |
| Viking 51 × IHO-4 (4)* | 0.43 | 0.57 | 0.43 | 0.21 | 0.39 | 0.12 |
| Viking 60 × IHO-2 (5)* | 0.54 | 0.10 | 0.48 | 0.14 | 0.79 | 0.13 |
| CCO-1 × IHO-1 (6) | 0.62 | 0.79 | 0.50 | 0.37 | 0.59 | 0.23 |
| CCO-1 × IHO-3 (7) | **0.00** | **0.22** | **0.00** | **0.21** | **0.00** | **0.37** |
| CCO-1 × IHO-4 (8) | 0.38 | 0.72 | 0.11 | 0.67 | 0.00 | 0.70 |
| CCO-2 × IHO-1 (9) | 0.79 | 0.29 | 0.64 | 0.67 | 0.48 | 0.93 |
| CCO-2 × IHO-2 (10) | 0.72 | 0.16 | 0.77 | 0.30 | 0.75 | 0.42 |
| CCO-2 × IHO-3 (11) | 0.45 | 0.60 | 0.55 | 0.40 | 0.58 | 0.59 |
| CCO-2 × IHO-4 (12) | 0.00 | 0.85 | 0.02 | 0.69 | 0.23 | 0.63 |
| **Mean** | **0.38** | **0.45** | **0.32** | **0.38** | **0.35** | **0.44** |

<sup>‡</sup>Three multivariate methods: partial least squares discriminant analysis (PLS-DA), support vector machine (SVM), and random forest (RF)
<sup>1</sup>False discovery rate (FDR) is the proportion of predicted haploid kernels that were actually hybrids
<sup>2</sup>False negative rate (FNR) is the proportion of true haploid kernels that were incorrectly predicted as hybrids
\*Population met $FDR \leq 0.25$ and $FNR \leq 0.50$ operational benchmarks under population-specific PLS-DA model
Bolded values indicate populations that passed the $FDR \leq 0.25$ and $FNR \leq 0.50$ operational benchmarks under the given multivariate general model. General models were trained on 11 populations using all haploid and hybrid kernels, while the models were evaluated on the 12<sup>th</sup> held-out population using hold-one-population-out cross-validation (HOPO-CV) with the test set resampled to the population-specific haploid induction rate (HIR), and including defectives in the test set

To evaluate whether restricting the training set to populations with stronger spectral discrimination would improve general model performance, an additional general model was trained and tested on only the 8 Viking and CCO-1 donor populations, excluding the four CCO-2 populations. Excluding CCO-2 populations from training did not improve classification accuracy for the remaining populations. The Viking + CCO-1 general SVM model showed slightly higher mean FDR (0.305 vs. 0.234) and virtually identical mean FNR (0.328 vs. 0.318) relative to the all-12 general SVM model when evaluated on the same 8 populations, suggesting that CCO-2 populations contribute a small amount of additional spectral information to the general model that improves classification accuracy in Viking and CCO-1 donor populations. The performance of the two models also suggests that the general SVM model was not negatively impacted by the inclusion of CCO-2 populations in the training set despite their poor individual classification accuracy.

The relationship between Δ oil and general SVM classification error rates was examined across all 12 populations (**Fig. 5**). A negative correlation was observed between Δ oil and both FDR (*r* = −0.49) and FNR (*r* = −0.34), with populations that showed larger oil content differences between haploid and hybrid kernels tending to have lower classification error rates under the general SVM model. CCO-1 × IHO-1 (6) was a noticeable outlier in the FDR relationship. Despite having the largest Δ oil of all 12 populations (2.54%), it showed the highest FDR among populations with Δ oil > 1.6% (FDR = 0.50). CCO-1 × IHO-1 (6) also had the second lowest HIR (4%) of all 12 populations. The negative trend between Δ oil and FNR was weak across populations, with most populations falling under the 0.50 operational benchmark.

**Fig. 5.**
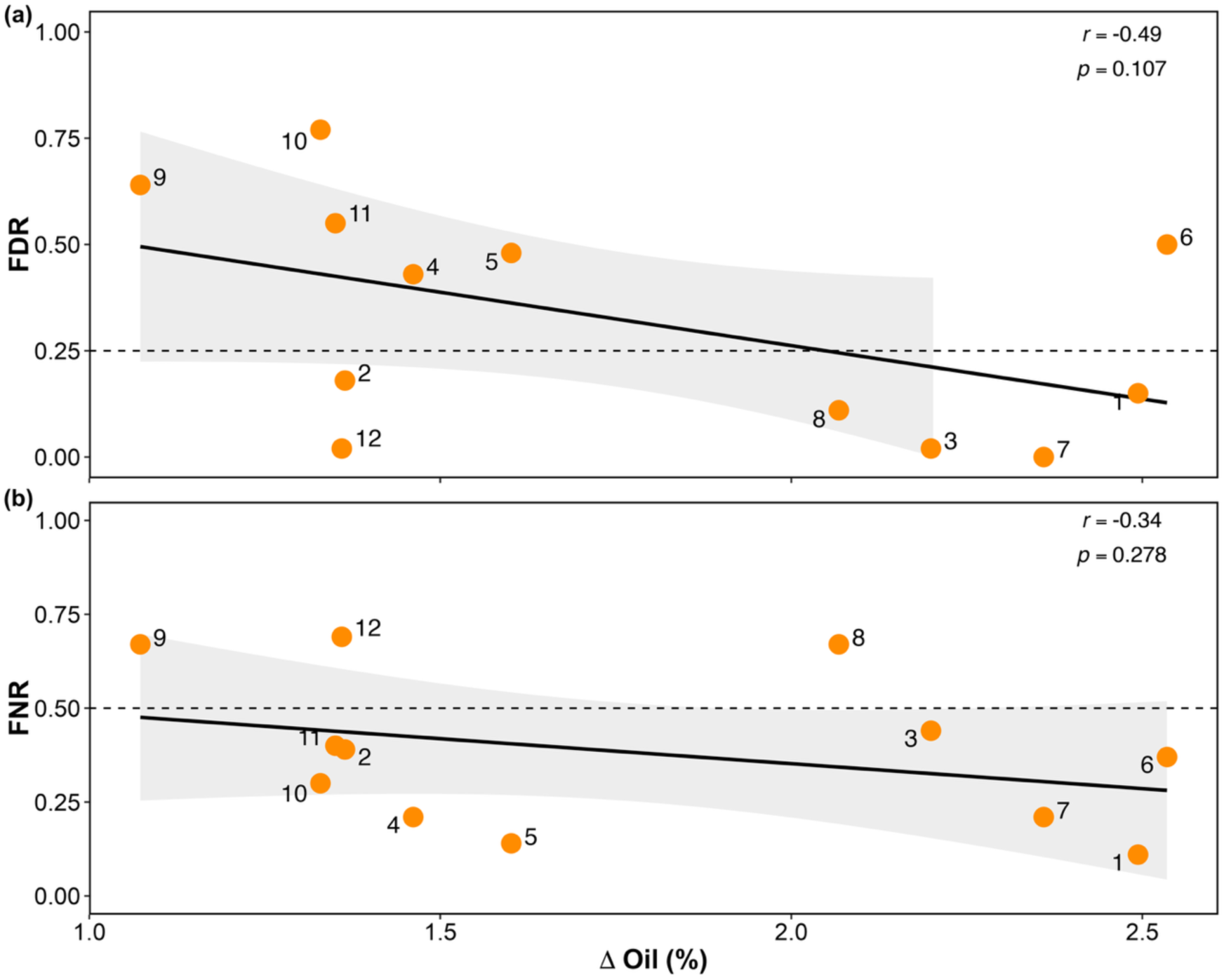
Relationship between Δ oil and the **(a)** false discovery rate (FDR) **(b)** false negative rate (FNR) from the general SVM model across 12 induction populations. Each point represents a population, with numbers corresponding to population IDs (1-12). Pearson correlation coefficients (*r*) and associated *p*-value are shown in each panel. Dashed horizontal lines indicate the operational benchmarks of FDR ≤ 0.25 (upper panel) and FNR ≤ 0.50 (lower panel). Solid lines represent least-squares linear regression fits, shaded areas representing 95% confidence intervals.

### 3.5. Multivariate spectral classification for defective kernel sorting

A secondary classification model was developed to remove defective kernels from the predicted haploid pool following the initial haploid sorting step. All three classification methods achieved low FDR and FNR across all 10 populations with defective kernels (**Supplementary Table 4**). Mean FDR across all populations was 0.014, 0.022, and 0.033 for PLS-DA, SVM, and RF, respectively, while mean FNR was 0.006, 0.015, and 0.015, respectively. No population showed FDR or FNR exceeding 0.10 under any method. PLS-DA achieved the lowest mean FDR (0.014) and FNR (0.006), indicating that nearly all non-defective kernels were correctly identified and retained in the predicted haploid pool. CCO-1 × IHO-3 (7) showed perfect classification under all three methods with FDR and FNR values equal to zero. The highest FDR observed under PLS-DA was 0.045 for Viking 51 × IHO-1 (1). Kernels were determined to be defective based on visual scoring but were not confirmed with a secondary method and may contain errors that could increase the prediction error.

## 4. Discussion

Accurate haploid identification is a critical step in DH generation in maize breeding, yet the dominant approach, manual classification based on the *R1-nj* kernel color marker, is subjective, labor-intensive, and unreliable in sweet corn backgrounds where anthocyanin accumulation is suppressed (Tracy 2001; Chaikam et al. 2015; reviewed by Dermail et al. 2024b). In this study, we evaluated an skNIR sorter as an automated, pigmentation-independent alternative for haploid classification across 12 induction populations comprising two field corn and two sweet corn donors crossed with four HOHIs. Two complementary sorting approaches were assessed: oil-based methods that exploit the relative difference in oil content between hybrid and haploid kernels (Δ oil) produced with HOHIs and multivariate methods that classify kernels using all wavelengths of the skNIR spectra (940-1640 nm) rather than a single predicted trait.

Visual *R1-nj*-based haploid sorting achieved acceptable accuracy across most field corn populations but was substantially less reliable in sweet corn backgrounds, where anthocyanin suppression approximately doubled sorting error rates relative to field corn, underscoring the need for pigmentation-independent approaches in those backgrounds. In this study, NIR-based haploid kernel sorting was achievable in a subset of the 12 induction populations evaluated, but performance varied substantially across populations regardless of the sorting approach applied. The general multivariate SVM model, applied without population-specific recalibration, achieved acceptable classification accuracy in four of 12 induction populations, outperforming oil-based methods, which met operational benchmarks in only two of 12 populations. Developing a separate calibrated model for each induction cross improved classification accuracy by meeting performance thresholds in only one additional population (5 of 12), indicating that the majority of unsortable populations reflect insufficient detectable differences between haploid and hybrid kernels rather than a limitation of the modeling approach. Two of the four IHO inducers drove sufficient Δ oil such that most, but not all, of the induction populations created with these inducers could be sorted. Continued development of HOHI lines capable of driving larger, more consistent Δ oil across a broader range of donor backgrounds would improve the general utility of the skNIR-based sorting system for any induction cross.

Excluding anthocyanins, the strongest chemical signature that distinguishes haploid from hybrid kernels is relative oil content (Melchinger et al. 2015a). However, oil content-based haploid sorting with the skNIR platform is constrained by the accuracy with which oil content can be predicted from NIR spectra. The PLS model implemented on the skNIR system used for oil content predictions achieved R² values of 0.54-0.60 across the three kernel classes against NMR-measured values (**Supplementary Fig. 2**). This suggests moderate predictive ability that introduces error into oil-based classification before any sorting threshold is applied. Classification methods based directly on NMR-based oil content estimates have achieved substantially higher sorting accuracy in previous studies (Liu et al. 2012; Melchinger et al. 2014, 2015a, b; Dong et al. 2014; Wang et al. 2016; Ge et al. 2020), and a data fusion approach combining NIR and NMR spectral data has also shown improved discrimination of haploid and hybrid kernels relative to either method alone (Ge et al. 2021). Despite this accuracy advantage, NMR systems are comparatively slower, processing kernels at approximately 4 seconds per kernel compared to 0.25 seconds per kernel for the skNIR sorter, and require larger, more expensive equipment less suited for high-throughput production environments (Wang et al. 2016; Armstrong 2006). Improving the accuracy of the PLS oil prediction model, for example by incorporating a broader diversity of kernel types and genetic backgrounds in the calibration set, would help increase the effectiveness of skNIR as a haploid sorting technology.

Oil-based sorting is the most straightforward approach when an induction population exhibits a bimodal oil content distribution, with haploid and hybrid kernels forming distinct, non-overlapping groups (Melchinger et al. 2013). However, none of the 12 induction populations in this study showed clear bimodality, and the optimal threshold for separating haploids from hybrids changed for each population. We therefore tested two general threshold-setting approaches: the lowest percentage method that classifies a fixed proportion of the lowest-oil kernels as haploid, and a dynamic threshold method that sets the haploid cutoff at a fixed distance below the population mean oil content. These methods could be generally applicable to any induction population created using a HOHI without recalibration of the instrument. Each method met operational benchmarks in only two of 12 populations and represent a pronounced FDR-FNR tradeoff: the dynamic threshold method produced a cleaner haploid pool (low FDR) but at the cost of greater haploid loss (high FNR) relative to the lowest percentage method. When oil content distributions overlap substantially between kernel classes, no fixed general threshold can simultaneously minimize both error types. Consistent with Gustin et al. (2020), who showed that skNIR-based haploid sorting is feasible when Δ oil exceeds approximately 1.7% using a conventional inducer, the HOHIs evaluated here approximately doubled mean Δ oil relative to that study (1.77% vs. 0.88%), supporting their value for improving oil-based sorting performance on the skNIR platform.

The three multivariate methods evaluated here avoid oil content predictions entirely by classifying kernels directly using the skNIR spectra. Multivariate classification methods outperformed oil-based sorting methods across all model configurations evaluated, likely because full-spectrum classification avoids the error introduced by intermediate oil content predictions and captures additional spectral information beyond oil accumulation alone (Gustin et al. 2020). Previous studies evaluating NIR spectroscopy coupled with multivariate classification methods have reported haploid identification accuracies of 90-100% in maize (Jones et al. 2012; Yu et al. 2018; Wang et al. 2018; Cui et al. 2019; He et al. 2022; Rodrigues Ribeiro et al. 2023; Kahrıman et al. 2026), but none of these studies included defective kernels in model evaluation to simulate realistic sorting operation. It was observed that excluding defective kernels from the test set substantially inflates apparent classification accuracy and can cause additional populations to appear sortable, suggesting the high accuracies reported in prior studies may overestimate performance under realistic induction population compositions. Among multivariate models tested, the general SVM model using HOPO-CV showed the best performance in distinguishing haploid and hybrid kernels during the initial haploid sorting step, which is consistent with previous findings in chemometric and image-based modeling for SVM-based classification (Xu et al. 2006; Kahrıman et al. 2023).

The effectiveness of haploid sorting is largely influenced by the genetic backgrounds of both the donor and haploid inducer used to generate haploid induction crosses. Kernel oil is primarily stored in the embryo, which reflects only the maternal genome in haploid kernels but both maternal and paternal contributions in hybrids (Rotarenco et al. 2007). In hybrid kernels, the paternal contribution from the haploid inducer can increase oil content through the xenia effect, which is amplified when high-oil inducers are used (Chen and Song 2003; Melchinger et al. 2013, 2014; Dong et al. 2014). However, some donors are resistant to HOHI-driven Δ oil inflation, resulting in greater overlap between kernel classes regardless of which inducer is used. The basis for this resistance is not understood, and no clear *a priori* signal for identifying resistant donors has yet been found. Donor oil content does not appear to predict responsiveness, as Melchinger et al. (2015b) found no relationship between the oil content of the donor and Δ oil. If the xenia effect on kernel oil content is heritable, the Δ oil produced by a given donor-inducer combination may instead be predictable through genomic prediction. Such predictions would be the most valuable for matching each donor to the inducer and sorting strategy best suited to it. Donors that respond strongly to HOHIs could be paired with HOHIs and sorted with the skNIR platform, while donors that retain reliable anthocyanin marker expression could be paired with conventional *R1-nj* inducers and sorted visually. Donors that are both resistant to HOHI-driven Δ oil inflation and that inhibit anthocyanin production, such as CCO-2, fall outside of both strategies and represent the central challenge for fully automating haploid sorting. Donor background accounted for 60.3% of the total variance in Δ oil across the 12 induction populations, accounting for nearly four times the variance attributable to the inducer (15.6%; **Supplementary Table 1**). Populations with Viking 51 and CCO-1 donor backgrounds generated the largest Δ oil and were the most amenable to sorting under both oil-based and multivariate methods, whereas CCO-2 populations consistently produced the lowest Δ oil and could not be accurately sorted under any method evaluated, regardless of inducer. The frequency of this donor’s resistance to HOHI-driven Δ oil inflation across germplasm is not well characterized, though Melchinger et al. (2015b) and Dong et al. (2014) each reported it in only a minority of field corn donors, suggesting that donor resistance may be relatively uncommon in field corn germplasm. Whether sweet corn backgrounds are more frequently resistant warrants further investigation.

Furthermore, it would be useful to investigate whether donors resistant to one HOHI would be responsive to a HOHI from a different genetic background. Inducer background contributed less to Δ oil variation than donor background, but the inducers driving the largest Δ oil also had the lowest HIR, consistent with Dong et al. (2014). Because low HIR raises the induction population size needed to recover haploids and can increase sorting error, this tradeoff poses a constraint on breeding HOHIs that combine large Δ oil with high HIR.

Low HIR can also arise from the donor; some donor genotypes are partially incompatible with a given inducer or unresponsive to induction, resulting in reduced seed yield and HIR, thereby increasing DH production costs (Trentin et al. 2022). From a plant breeding standpoint, these donor-side genetic effects are a major obstacle for the practical implementation of DH technology (Dermail et al. 2024a). Donor selection is driven by breeding objectives that determine the agronomic performance of the resulting inbred lines and hybrids. Imposing selection pressure on HOHI responsiveness or HIR would constrain germplasm choice without any downstream benefit to line quality. The appropriate response to donor-dependent variation in Δ oil is therefore to develop HOHI lines that drive sufficient Δ oil across a broader range of donor backgrounds, rather than to restrict which donors enter the DH pipeline.

In addition to haploid and hybrid kernels, the haploid induction process introduces non-viable or defective kernels arising from embryo abortion, endosperm abortion, or incomplete kernel development (Zhao et al. 2013; Qu et al. 2020). Qu et al. (2020) found substantial overlap between QTL controlling kernel abortion and QTL for HIR, suggesting that defective kernel production may be an inherent feature of the *in vivo* haploid induction process. Effective handling of defective kernels is therefore an important consideration in the development of any haploid sorting system. Because defective kernels do not germinate, their misclassification as haploid wastes pre-germination resources at the planting and greenhouse stage but has no effect on the genetic composition or quality of the resulting DH population. We defined FDR and FNR using haploid and hybrid kernels only, as confusion between those two classes represents the only sorting error with direct consequences for DH production outcomes. Defective kernels were nonetheless retained in the test set to ensure that model performance estimates reflected the realistic composition of induction populations encountered in practice.

The predicted oil content distributions of defective and haploid kernels overlapped considerably, precluding accurate classification of these kernel classes using oil-based methods alone. The overlap in oil content is likely because disrupted or incomplete kernel development would be expected to reduce oil accumulation relative to hybrid kernels (Zhao et al. 2013; Liu et al. 2018). In contrast, the NIR spectral signatures of defective kernels were sufficiently distinct to support accurate classification by a dedicated defective kernel sorting model based on multivariate methods. Using a binary classification framework sorting non-defective vs. defective kernels, all three multivariate methods achieved FDR and FNR below 0.10 across all 10 populations containing defective kernels (**Supplementary Table 4**). Operationally, the most efficient approach is to apply this defective kernel cleaning step to the predicted haploid pool following the initial haploid sorting step, rather than running the entire induction population through a defective cleaning step prior to haploid sorting.

This study demonstrates that the skNIR sorter is a viable platform for rapid, objective haploid kernel sorting in maize DH production programs, including in sweet corn backgrounds where visual *R1-nj*-based sorting is unreliable due to suppression or inhibition of kernel color marker expression. Both oil-based and spectra-based multivariate approaches were able to accurately sort haploid kernels in a subset of the induction populations evaluated, with the general SVM model achieving the best overall classification accuracy among the deployable sorting approaches. Sorting accuracy was primarily influenced by donor genotype through its effect on Δ oil, and populations with insufficient Δ oil or very low HIR remained challenging for all methods evaluated. The strong performance of the dedicated defective kernel sorting model supports a two-step NIR-based sorting workflow for operational DH production.

Several avenues exist for improving the accuracy and generalizability of skNIR-based haploid sorting. The diversity of donor backgrounds included in general model training sets should be expanded to ensure more complete representation of the germplasm encountered in practical DH production programs, which would be expected to improve both classification accuracy and the range of populations for which general models are effective. In parallel, developing HOHI lines that drive consistently large Δ oil and high HIR across diverse donor genetic backgrounds would address the primary challenge for unrestricted deployment. By improving classification accuracy and throughput at the stage of haploid identification, the skNIR sorter has the potential to enhance DH production efficiency and reduce the cost per line, facilitating broader adoption of DH technology in maize breeding programs.

## Author contribution statement

Conceptualization: AMS, MR, UKF, TL, JLG, JH; Methodology: SS, JLG, AMS, MR, UKF, TL, JH; Formal analysis and investigation: SS, JLG; Writing - original draft preparation: SS, JLG, JH; Writing - review and editing: SS, JLG, UKF, AMS, TL, MR, JH; Funding acquisition: AMS, MR; Resources: JLG, UKF, TL, MR, JH; Supervision: TL, MR, JH

## Funding

This work was supported by the United States Department of Agriculture NIFA SCRI grants, award numbers 2018-51181-28419 and 2022-51181-38333.

## Acknowledgements

We thank Paul Armstrong for designing, constructing, and maintaining the skNIR sorter instrument and its software, implementing multivariate models on the instrument, and performing the NMR analysis. This research used in part resources on the Palmetto Cluster at Clemson University under National Science Foundation awards MRI 1228312, II NEW 1405767, MRI 1725573, and MRI 2018069. The views expressed in this article do not necessarily represent the views of NSF or the United States government.

## Declaration of competing interests

The authors declare that the research was conducted without any commercial or financial relationships that could be construed as a potential conflict of interest. Mention of trade names or commercial products in this publication is solely for the purpose of providing specific information and does not imply recommendation or endorsement by the U.S. Department of Agriculture. USDA is an equal opportunity provider and employer.

## Data availability

The datasets generated and analyzed in this study are available in the Dryad repository, DOI: 10.5061/dryad.cfxpnvxp1. Analysis R codes are available via GitHub, https://github.com/HershbergerLab/SweetCAP_skNIR_Haploid_Discriminant_Analysis.

## Notes

https://doi.org/10.5061/dryad.cfxpnvxp1

### Competing Interest Statement

The authors have declared no competing interest.

