## Supplementary for "Single-kernel near-infrared spectroscopy enables haploid kernel sorting in field and sweet corn using high-oil haploid inducers across diverse donor-inducer combinations"

### Supplementary Tables

**Supplementary Table 1** Two-way analysis of variance (ANOVA) estimation of the effect size ($\eta^2$) of induction population donors and inducers on delta oil (Δ oil) and haploid induction rate (HIR) in 12 induction populations

*Δ oil*

| Source | SS | df | F | *p*-value | $\eta^2$ | $\eta^2$ (%) |
|---|---|---|---|---|---|---|
| Donor | 1.6503 | 3 | 4.160 | 0.079 | 0.603 | 60.3% |
| Inducer | 0.4269 | 3 | 1.076 | 0.438 | 0.156 | 15.6% |
| Residual | 0.6611 | 5 | — | — | 0.241 | 24.1% |

*HIR*

| Source | SS | df | F | *p*-value | $\eta^2$ | $\eta^2$ (%) |
|---|---|---|---|---|---|---|
| Donor | 0.0044 | 3 | 7.604 | 0.026 | 0.540 | 54.0% |
| Inducer | 0.0028 | 3 | 4.814 | 0.062 | 0.342 | 34.2% |
| Residual | 0.0010 | 5 | — | — | 0.118 | 11.8% |

Type III SS. $\eta^2 = SS_{effect}/ SS_{total}$; $df_{error} = 5$

**Supplementary Table 2** Least squares (LS) means, 95% confidence interval (CI) and pairwise comparisons for donor and inducer effects on delta oil (Δ oil) and haploid induction rate (HIR)

| Donor | Δ oil LS mean (%) | SE | 95% CI | Group | HIR LS mean (%) | SE | 95% CI | Group |
|---|---|---|---|---|---|---|---|---|
| CCO-2 | 1.28 | 0.182 | 0.59 – 1.97 | a | 9.25 | 0.70 | 6.60 – 11.90 | ab |
| Viking 60* | 1.83 | 0.426 | 0.21 – 3.46 | a | 8.88 | 1.63 | 2.65 – 15.10 | ab |
| Viking 51 | 1.88 | 0.182 | 1.19 – 2.57 | a | 10.50 | 0.70 | 7.85 – 13.15 | b |
| CCO-1 | 2.24 | 0.223 | 1.40 – 3.09 | a | 5.38 | 0.85 | 2.12 – 8.63 | a |

| Inducer | Δ oil LS mean (%) | SE | 95% CI | Group | HIR LS mean (%) | SE | 95% CI | Group |
|---|---|---|---|---|---|---|---|---|
| IHO-2 | 1.58 | 0.223 | 0.73 – 2.42 | a | 9.63 | 0.85 | 6.37 – 12.88 | a |
| IHO-4 | 1.64 | 0.239 | 0.73 – 2.55 | a | 10.46 | 0.91 | 6.97 – 13.94 | a |
| IHO-3 | 1.98 | 0.239 | 1.07 – 2.89 | a | 7.13 | 0.91 | 3.64 – 10.61 | a |
| IHO-1 | 2.04 | 0.239 | 1.13 – 2.95 | a | 6.79 | 0.91 | 3.31 – 10.28 | a |

Least squares (LS) means estimated from main-effects model (Type III SS). Pairwise comparisons adjusted using Bonferroni correction (df error = 5). Means sharing the same letter are not significantly different at $\alpha = 0.05$.
*Viking 60 standard error (SE) is larger due to fewer combinations in the dataset.

**Supplementary Table 3** Classification error rates for 12 induction populations at the per-population optimal, lowest percentage (14%) and dynamic (offset = 2.0) oil content classification thresholds

| **Population** | **HIR (%)[3]** | **Per-population Optimal** | | **Lowest Percentage (14%)** | | **Dynamic Threshold (offset = 2.0)** | |
|---|---|---|---|---|---|---|---|
| | | **FDR[1]** | **FNR[2]** | **FDR[1]** | **FNR[2]** | **FDR[1]** | **FNR[2]** |
| Viking 51 × IHO-1 (1)* | 8 | **0.19** | **0.00** | **0.20** | **0.36** | 0.21 | 0.53 |
| Viking 51 × IHO-2 (2) | 13 | 0.31 | 0.49 | 0.38 | 0.45 | 0.10 | 0.91 |
| Viking 51 × IHO-3 (3)* | 10 | **0.23** | **0.01** | **0.25** | **0.03** | **0.10** | **0.46** |
| Viking 51 × IHO-4 (4)* | 11 | **0.09** | **0.46** | 0.14 | 0.67 | 0.00 | 0.99 |
| Viking 60 × IHO-2 (5) | 10 | 0.38 | 0.42 | 0.40 | 0.60 | 0.43 | 0.91 |
| CCO-1 × IHO-1 (6) | 4 | 0.33 | 0.40 | 0.61 | 0.06 | 0.39 | 0.46 |
| CCO-1 × IHO-3 (7)* | 3 | **0.00** | **0.12** | 0.79 | 0.11 | **0.00** | **0.19** |
| CCO-1 × IHO-4 (8)* | 8 | **0.25** | **0.28** | 0.38 | 0.19 | 0.38 | 0.62 |
| CCO-2 × IHO-1 (9) | 8 | 0.64 | 0.40 | 0.71 | 0.55 | 0.00 | 0.97 |
| CCO-2 × IHO-2 (10) | 9 | 0.55 | 0.44 | 0.65 | 0.59 | 0.60 | 0.85 |
| CCO-2 × IHO-3 (11) | 8 | 0.64 | 0.43 | 0.70 | 0.56 | 0.79 | 0.95 |
| CCO-2 × IHO-4 (12) | 12 | 0.34 | 0.47 | 0.39 | 0.46 | 0.00 | 0.87 |
| **Mean** | **9** | **0.33** | **0.33** | **0.47** | **0.39** | **0.25** | **0.73** |

[1]False discovery rate
[2]False negative rate
[3]Haploid induction rate
*Population met FDR ≤ 0.25 and FNR ≤ 0.50 operational benchmarks under the per-population optimal threshold. Bolded values indicate populations meeting the operational benchmarks.

**Supplementary Table 4** General model defective kernel classification error rates for three multivariate methods across 10 induction populations that contain defective kernels

| Population* | PLS-DA‡ | | SVM‡ | | RF‡ | |
|---|---|---|---|---|---|---|
| | FDR[1] | FNR[2] | FDR[1] | FNR[2] | FDR[1] | FNR[2] |
| Viking 51 × IHO-1 (1) | 0.045 | 0.003 | 0.045 | 0.003 | 0.045 | 0.003 |
| Viking 51 × IHO-2 (2) | 0.021 | 0.000 | 0.029 | 0.000 | 0.047 | 0.000 |
| Viking 51 × IHO-4 (4) | 0.042 | 0.023 | 0.048 | 0.010 | 0.057 | 0.023 |
| Viking 60 × IHO-2 (5) | 0.000 | 0.033 | 0.016 | 0.020 | 0.000 | 0.022 |
| CCO-1 × IHO-1 (6) | 0.031 | 0.000 | 0.047 | 0.066 | 0.076 | 0.032 |
| CCO-1 × IHO-3 (7) | 0.000 | 0.000 | 0.000 | 0.000 | 0.000 | 0.000 |
| CCO-1 × IHO-4 (8) | 0.000 | 0.000 | 0.000 | 0.027 | 0.029 | 0.019 |
| CCO-2 × IHO-2 (10) | 0.000 | 0.000 | 0.000 | 0.027 | 0.010 | 0.027 |
| CCO-2 × IHO-3 (11) | 0.000 | 0.000 | 0.020 | 0.000 | 0.041 | 0.024 |
| CCO-2 × IHO-4 (12) | 0.000 | 0.000 | 0.016 | 0.000 | 0.022 | 0.000 |
| **Mean (n=10)** | **0.014** | **0.006** | **0.022** | **0.015** | **0.033** | **0.015** |

‡Three multivariate methods: partial least squares discriminant analysis (PLS-DA), support vector machine (SVM), and random forest (RF)
[1]False discovery rate
[2]False negative rate
***Populations 3 and 9 contained no defective kernels

**Supplementary Figures**

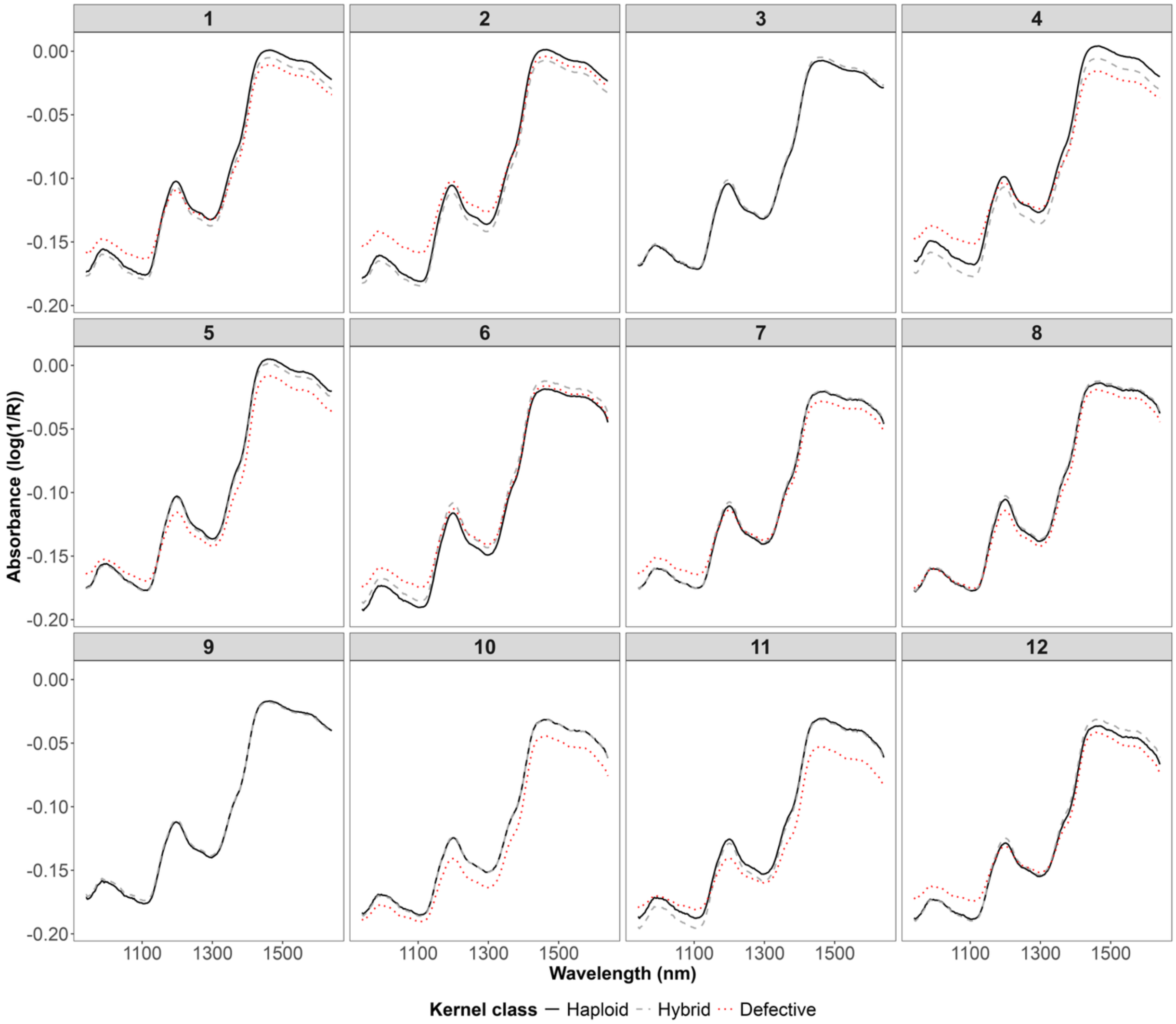


**Supplementary Fig. 1** Mean absorbance of the raw near-infrared (NIR) spectra of haploid, hybrid and defective kernels for each induction population [Viking 51 × IHO-1 (**1**), Viking 51 × IHO-2 (**2**), Viking 51 × IHO-3 (**3**), Viking 51 × IHO-4 (**4**), Viking 60 × IHO-2 (**5**), CCO-1 × IHO-1 (**6**), CCO-1 × IHO-3 (**7**), CCO-1 × IHO-4 (**8**), CCO-2 × IHO-2 (**9**), CCO-2 × IHO-2 (**10**), CCO-2 × IHO-3 (**11**), CCO-2 × IHO-4 (**12**)]. Absorbance is reported as log(1/Reflectance) for the NIR wavelength range of 940 to 1640 nm using the single-kernel NIR reflectance spectroscopy sorter

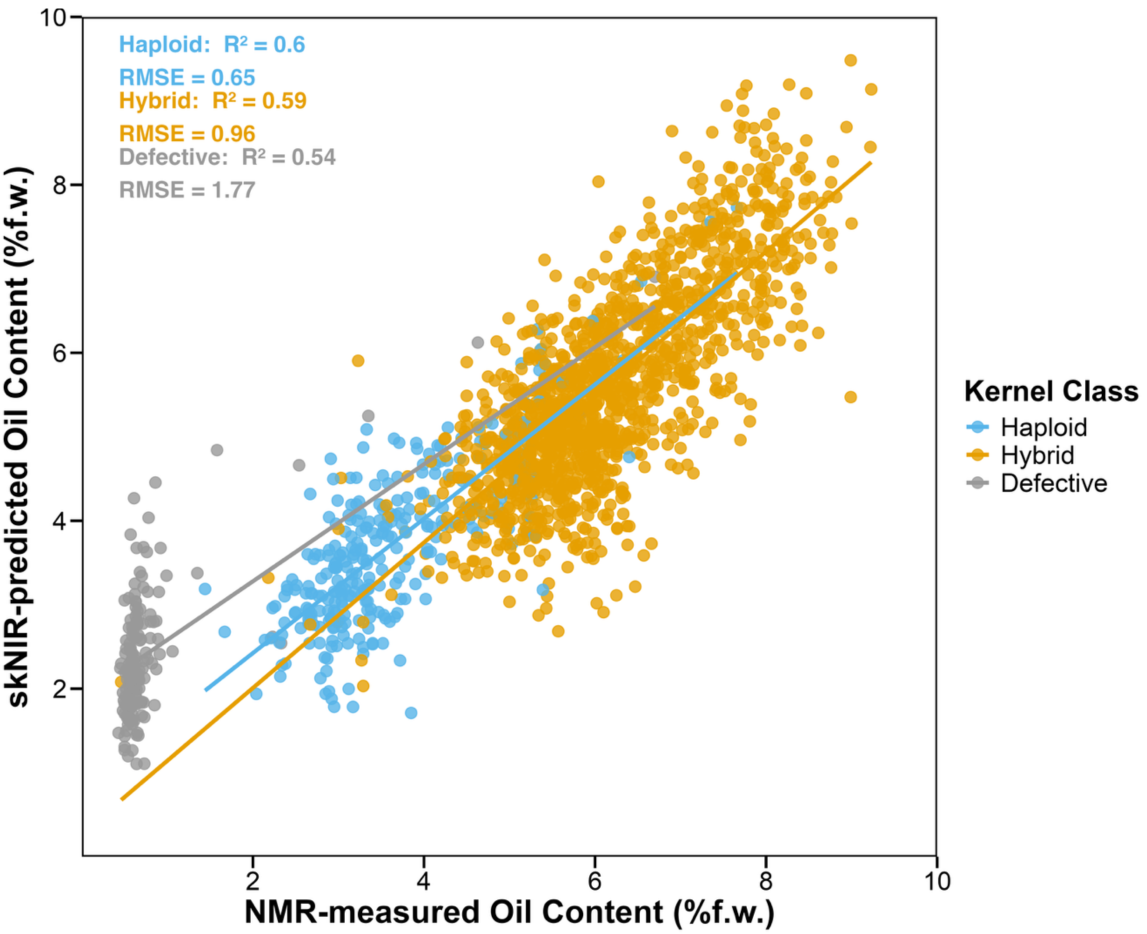


**Supplementary Fig. 2** Comparison of single-kernel near-infrared reflectance spectroscopy (skNIR)-predicted oil content (%) with nuclear magnetic resonance (NMR)-measured oil content (%) of the three kernel classes across 9 induction populations
